# A yeast BiFC-seq method for genome-wide interactome mapping

**DOI:** 10.1101/2020.06.16.154146

**Authors:** Limin Shang, Yuehui Zhang, Yuchen Liu, Chaozhi Jin, Yanzhi Yuan, Chunyan Tian, Ming Ni, Xiaochen Bo, Li Zhang, Dong Li, Fuchu He, Jian Wang

**Author notes:** Correspondence should be addressed to F.H. and J.W. Corresponding authors. (Fuchu He), (Jian Wang).

## Abstract

Genome-wide physical protein-protein interaction (PPI) mapping remains a major challenge for current technologies. Here, we report a high-efficiency yeast bimolecular fluorescence complementation method coupled with next-generation DNA sequencing (BiFC-seq) for interactome mapping. We applied this technology to systematically investigate an intraviral network of Ebola virus (EBOV). Two-thirds (9/13) of known interactions of EBOV were recaptured and five novel interactions were discovered. Next, we used BiFC-seq method to map the interactome of the tumor protein p53. We identified 97 interactors of p53 with more than three quarters are novel. Furthermore, in more complex background, we screened potential interactors by pooling two BiFC-libraries together, and revealing a network of 229 interactions among 205 proteins. These results show that BiFC-seq is a highly sensitive, rapid and economical method in genome-wide interactome mapping.

## Introduction

Genome-wide yeast two hybrid (Y2H) screening and affinity-purification coupled with mass spectrometry (AP-MS) have been extensively used for mapping the interactomes of various species [1]-[4]. However, only limited coverage was obtained for the interactome of most organisms [5],[6], which dues to low sensitivity, labor intensive and high cost of the technologies.

The bimolecular fluorescence complementation (BiFC) assay is a powerful tool to investigate the binary protein-protein interactions. A Venus-based BiFC method has been used in large-scale interactome mapping of telomere signaling and SUMO [7]8] which specifically detected transient or weak interactions that cannot be obtained by Y2H or AP-MS. However, there is high background fluorescence intensity, which leads to more artificial interactions [9].

To overcome these limitations, we developed a BiFC-seq method, which combines the yEGFP-BiFC with NGS technology, expanding the BiFC application in genome-wide interactome screening. Firstly, we have used yEGFP-BiFC method to depict a small-scale intraviral PPI network of EBOV among its 9 encoding proteins, which contains 14 interactions among 6 proteins. Then, we applied BiFC-seq in high-throughput assay to screen p53 interactors from human universal library and revealed 97 p53 interactors with 21 reported in literature. Finally, we carried out genome-wide interactome screening by pooling two tagged libraries together using BiFC-seq, generating an interaction network consists of 229 interactions with 12 reported in the BioGrid database.

## Methods

### Data availability

The NGS data were uploaded to iProx database (http://www.iprox.org/). The sequencing data of p53 interactors are accessible with https://www.iprox.org/page/PSV023.html;?url=1592296064595kJiq and Password: yjJp. The sequencing data of genome-wide PPI are accessible with https://www.iprox.org/page/PSV023.html;?url=1592296389579vYEx and Password: U7Gw

### Construction of the yEGFP-BiFC vectors

The yEGFP-BiFC constructs were generated using the pDBleu and pPC86 vectors (Invitrogen, Grand Island, NY) as templates according to standard molecular techniques. Briefly, the coding regions of the DNA activation or binding domains were removed, and the DNA fragments coding for the N or C-terminal fluorescent proteins and a linker coding a 15 amino acid spacer sequence (GGGGS)3 were inserted. The remaining vectors were constructed with these backbone vectors using standard molecular techniques.

### Construction of the cDNA library

The cDNA library was constructed using Gateway recombination technology. Human universal reference total RNA (Catalog No. 636538, Clontech, Mountain View, CA) was used as a template to synthesize cDNA by reverse transcription. The cDNA was ligated to adaptors with the *att*B1 site. To construct the Gateway Entry vectors, pDONR222 (Catalog No. 636538, Invitrogen, Grand Island, NY) was mixed with purified cDNA and the BP Clonase™ II enzyme. The reactions were incubated at 25°C overnight and treated with Proteinase K at 37°C for 10 minutes to terminate the reaction. The reaction products were transformed into *E. coli* DH10B competent cells, colonies were grown in LB medium with kanamycin selection, and plasmids were extracted with a Plasmid Mini Kit (Catalog No. 12125, Qiagen, Hilden, Germany). To construct the cDNA library, pDONR222 entry vectors were mixed with pPC86-YN157-CCDB vectors and the Gateway LR Clonase II enzyme. The reactions were incubated at 25°C overnight and treated with Proteinase K at 37°C for 10 minutes to terminate the reaction. The reaction products were transformed into *E. coli* DH10B competent cells, colonies were grown in LB medium with ampicillin selection, and plasmids were extracted and stored at −80°C.

### yEGFP-BiFC assay

The yeast strain *S. cerevisiae* AH109 (genotypes *MAT*a, *trp*1-901, *leu*2-3, 112, *ura*3-52, *his*3-200, *gal*4△, *gal*80△, *LYS*2:*GAL*1_*UAS*_-*GAL*1_*TATA*_-*HIS*3, *GAL*2_*UAS*_-*GAL*2_*TATA*_-*ADE*2, *URA*3:*MEL*1_*UAS*_-*MEL*1_*TATA*_-*lacZ*) (Clontech, Mountain View, CA) was used in the BiFC screening. A small-scale sequential transformation procedure was performed using protocols from the manufacturer. After transformation, the yeast cells were resuspended in 10 ml of liquid SD without tryptophan and leucine (SD-2) medium, cultured in a 30°C shaker for 24 hours and incubated at 4°C for an additional 48 hours for fluorophore maturation. The yeast cells were collected by centrifugation at 7000 rpm for 30 s and resuspended with PBS. The yeast cells with reconstituted yEGFP were excited with a 488 nm laser and collected through a FACSAria III (BD Biosciences, Franklin Lakes, NJ) 530/30 nm bandpass filter. The sorted yeast cells were collected with PBS, spread onto SD-2 plates and cultured in a 30°C incubator for 2 days until yeast colonies reached 2-3 mm in diameter. Fluorescence at 488 nm excitation and 519 nm emission were observed under an inverted fluorescence microscope (Olympus, Tokyo, Japan).

### Yeast colony PCR

The yeast colonies were picked and digested with 20 μl 0.02 N NaOH. After 5 minutes of boiling, 2 μl of the solution was used as a template to amplify the inserted cDNA with the forward primer (5’-ATGGCTGACAAACAAAGATCTGGT-3’) and the reverse primer (5’-GCGGCCGCACCACTTTGTACAAGAAA-3’). The reaction buffers were mixed and aliquoted by a Biomek FX Laboratory Automation Workstation (Beckman Coulter, Brea, CA) into 384-well PCR plates. The PCR reactions were performed with I-5™ 2×High-Fidelity Master Mix (MCLAB, TSINGKE Biological Technology, China) according to the manufacturer’s instructions. For each amplification, 3 μl of the PCR products was mixed and purified using the QIAquick PCR Purification Kit (Catalog No. 28104, Qiagen, Hilden, Germany).

### NGS and BiFC-seq workflow

The purified PCR products were sheared with a Covaris S220 system (Applied Biosystems, Carlsbad, CA). To prepare the libraries, the sheared PCR products were processed using a Nextera XT DNA Sample Preparation Kit according to the manufacturer’s instructions (Illumina, San Diego, CA). All the libraries were sequenced at a 2×150 bp read length on an Illumina HiSeq 2000 platform. The images generated by the HiSeq 2000 were converted to raw reads using base calling (CASAVA version 1.8). After trimming the adaptor sequences from the raw reads, the reads with ≥10% unidentified bases (Ns) and ≥50% low-quality bases (PHRED quality scores ≤5) were also removed to implement quality control. The filtered reads were mapped to the GRCh38 genome (Human GENCODE version 22) by TopHat [10]. The raw counts of the sequencing reads for each transcript were calculated by the HTSeq Python package [11]. DESeq2 was then used for different expression analysis between p53 and control groups [12]. A Gene Ontology analysis was then performed with the clusterProfiler R package (http://www.bioconductor.org/packages/devel/bioc/html/ClusterProfiler.html).

### Co-IP assay

HEK293 cells were cultured in Dulbecco’s modified Eagle’s medium (Invitrogen, Grand Island, NY) with 10% (v/v) fetal bovine serum (FBS). After 48 hours of transfection, the HEK293 cells with Flag-p53 and Myc-interactors were harvested and lysed in HEPES lysis buffer [0.5% NP-40, 150 mM NaCl, 1% (v/v) Tween 20, 50 mM Tris–HCl (pH 7.5), 0.1% 1 M DTT, and a protease inhibitor cocktail]. The immunoprecipitations were performed using anti-Myc antibodies (Sigma-Aldrich, St. Louis, MO) and protein A/G-agarose (ThermoFisher, San Jose, CA) at 4°C. The lysates and immunoprecipitates were detected by the anti-Flag antibody, which was followed by detection with the ECL substrate (ThermoFisher, San Jose, CA).

## Results and discussion

### Development of yEGFP-based BiFC

A codon-optimized yeast enhanced green fluorescent protein (yEGFP) [13] that was split into nonfluorescent halves at the 157 aa position to yield YN157 (1-157 aa) and YC157 (158-238 aa) fragments. The intact yEGFP had a strong fluorescence signal in yeast, whereas the split yEGFP produced undetectable fluorescence (Figure S1). To verify that this approach could be used for detecting PPIs, the fragments of yEGFP linked by (GGGGS)3 peptide sequence were fused to the N-terminal of b-Jun, b-Fos and ΔbFos as positive and negative indicators (**Figure 1A**). The co-transfection of YN157-bJun and YC157-bFos led to the reassembly of yEGFP and showed a strong fluorescence signal. However, the combination of YN157-bJun and YC157-ΔbFos showed slight background fluorescence (**Figure 1B**). The calculated signal to noise ratio (S/N) of the yEGFP-BiFC method was 33:1 for all the cells and 4:1 for the fluorescent-only cells, which allows for high-sensitivity detection of PPIs. The percentage of fluorescent cells in the positive group was 8-fold higher than that in the negative group under identical transfection conditions (**Figure 1C**). As expected, the fluorescent positive cell count was 16.5% by FACS, which was approximately 33-fold higher than the negative control (**Figure 1D**). The different fusion patterns of interacting proteins with the yEGFP fragments potentially affected the proximity of the split fluorescent protein fragments. Thus, we constructed b-Jun and b-Fos N-terminal or C-terminal fused proteins and tested these combinations in yeast (Figure 1a). We found that only the N-terminal fusion of YN157-bJun and the N- or C-terminal fusions of YC157-bFos and bFos-YC157 produced strong BiFC signals, whereas the other combinations produced slight fluorescent signals (Figure 1C). These results suggest that the yEGFP-based BiFC system is suitable for PPI screening in yeast and indicate that variation in the fusing of yEGFP fragments to targets can potentially affect screening results.

**Figure 1.**
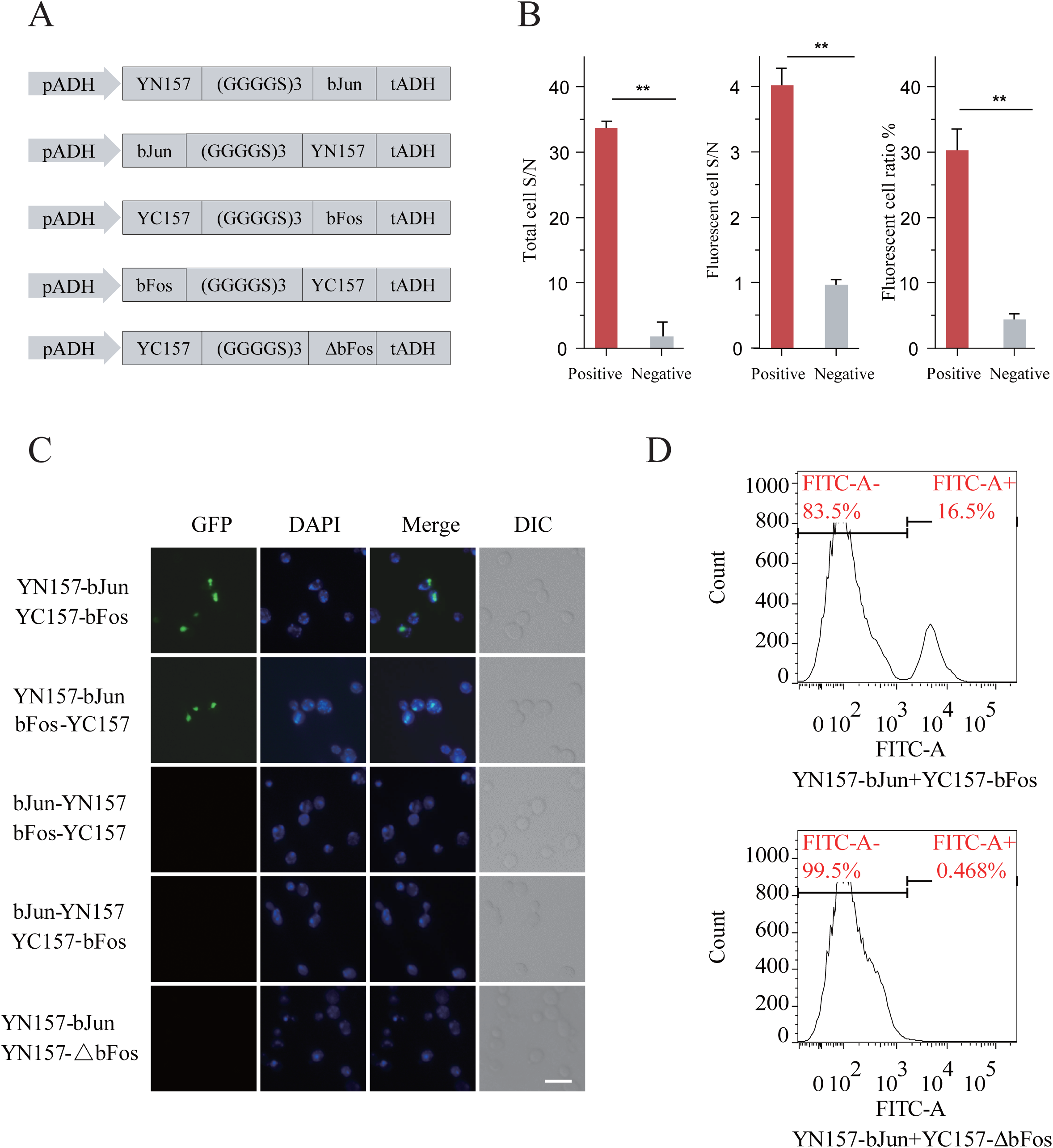
Development of yEGFP-based BiFC. (A) Diagram of different fusion patterns of yEGFP splited fragments of control groups. (B) Yeast cells cotransformed with control groups as indicated were detected under fluorescence microscope, the nuclei were labeled with DAPI. DIC: differential interference contrast microscopy scale bar: 12.5 μm. (C) Quantitative results of the fluorescence intensity of the yeast fluorescent cells. Total cell signal to noise ratio (S/N): 200 transformed yeast cells from positive and negative groups were randomly picked up, fluorescent intensities were analyzed by ImageJ and sumed up from each group to calculate Total S/N. Fluorescent cell S/N: the fluorescent intensities of 200 fluorescent cells from positive and negative groups were sumed up from each group to calculate Fluorescent cell S/N; and fluorescent cell ratio: the percentage of fluorescence-positive cells out of the total cells. Data are presented as means ± SD (n=3). Error bars: s.d. \*\**P*<0.01, Student’s t-test. (D) Approximately 20,000 yeast cells from positive and negative controls were analyzed by flow cytometry, the fluorescence-positive cell ratios were 16.5% and 0.5% respectively.

### An intraviral PPI network of Ebola virus by yEGFP-BiFC

The recent West African epidemic of EBOV disease caused more than 11,000 deaths owing to its high infectivity and lethality. Although tremendous efforts have been made to the development of EBOV therapeutics, an effective treatment remains elusive. The researches of pathogenic mechanisms of EBOV were mainly based on the pathogen-host PPI features [14],[15]. However, the intraviral PPI network is still unclear. Therapeutic strategy targeting interactions among internal EBOV proteins can be promising. Thus, we carried out a comprehensive analysis of interactions of EBOV proteins with each other using yEGFP-based BiFC. The EBOV genome encodes nine proteins: nucleoprotein (NP), viral protein (VP24, VP30, VP35 and VP40), glycoprotein (GP) and polymerase (L). The GP gene encodes for another two proteins (sGP and GP2) through RNA editing [16].

To exhaustively explore the PPI networks of 9 encoding proteins of EBOV, we fused each protein with YN157 or YC157 fragment, all in both orientations, yielding 36 fluorescent fragment tagged proteins. After validation of their expressions in yeast cells (Figure S2), these tagged proteins were pairwisely cotransformed into yeast cells in an 18×18 matrix format in triplicate to screen intraviral PPIs of EBOV, bJun and bFos/ΔbFos were used as controls. Transformed cells were analyzed by flow cytometry and fluorescence microscope (Figure S3). The ratio of fluorescent cells in each tested group was compared with the negative control (bJun and ΔbFos), and those with significant difference (*P*<0.05) and the log_2_ ratio (test/control) greater than 2 were regarded as authentic PPIs. Finally, 51 interactions among 19 tagged proteins were obtained (**Figure 2A**). After removing redundancy, the dataset contains 14 interactions among 6 EBOV proteins, of which 9 interactions were reported (**Figure 2B**). The interaction patterns of different orientations between fluorescent fragment tags and EBOV proteins varied greatly by BiFC method. The VP35-NP interaction can be detected by all pairwise combinations of tagged VP35 and NP proteins, whereas the VP24-GP2 interaction can only be detected by VP24-YC157 and GP2-YN157 combination (Figure S3), indicating the advantage of different fusion combinations in overcoming steric hindrance to detect PPIs by BiFC technology. Furthermore, by overexpressing Myc- and Flag-tagged EBOV proteins, we confirmed 10 of 14 interactions (71.4%) by co-immunoprecipitation assay (co-IP) in mammalian cells (**Figure 2C**). Our new findings are consistent with their biological roles, for example, the interactions of VP24-VP30 and VP24-GP2 suggest the role of VP24 in ribonucleoprotein complex (RNP) assembly and release process of GP2 out of the virus. The interactions of VP40 with VP24 and VP30 also confirmed that the VP40 involves in the process of releasing of viral particles from the host cells. By integrating our EBOV-BiFC protein networks with literature-curated datasets, a global intraviral EBOV protein networks containing 18 interactions among 7 proteins was obtained (Figure 2B) [17]-[23]. The comprehensive description of interplay among EBOV proteins provide a basis for studying protein functions in the aspect of viral replication and particle secretion as well as for advancing novel therapeutic approaches.

**Figure 2.**
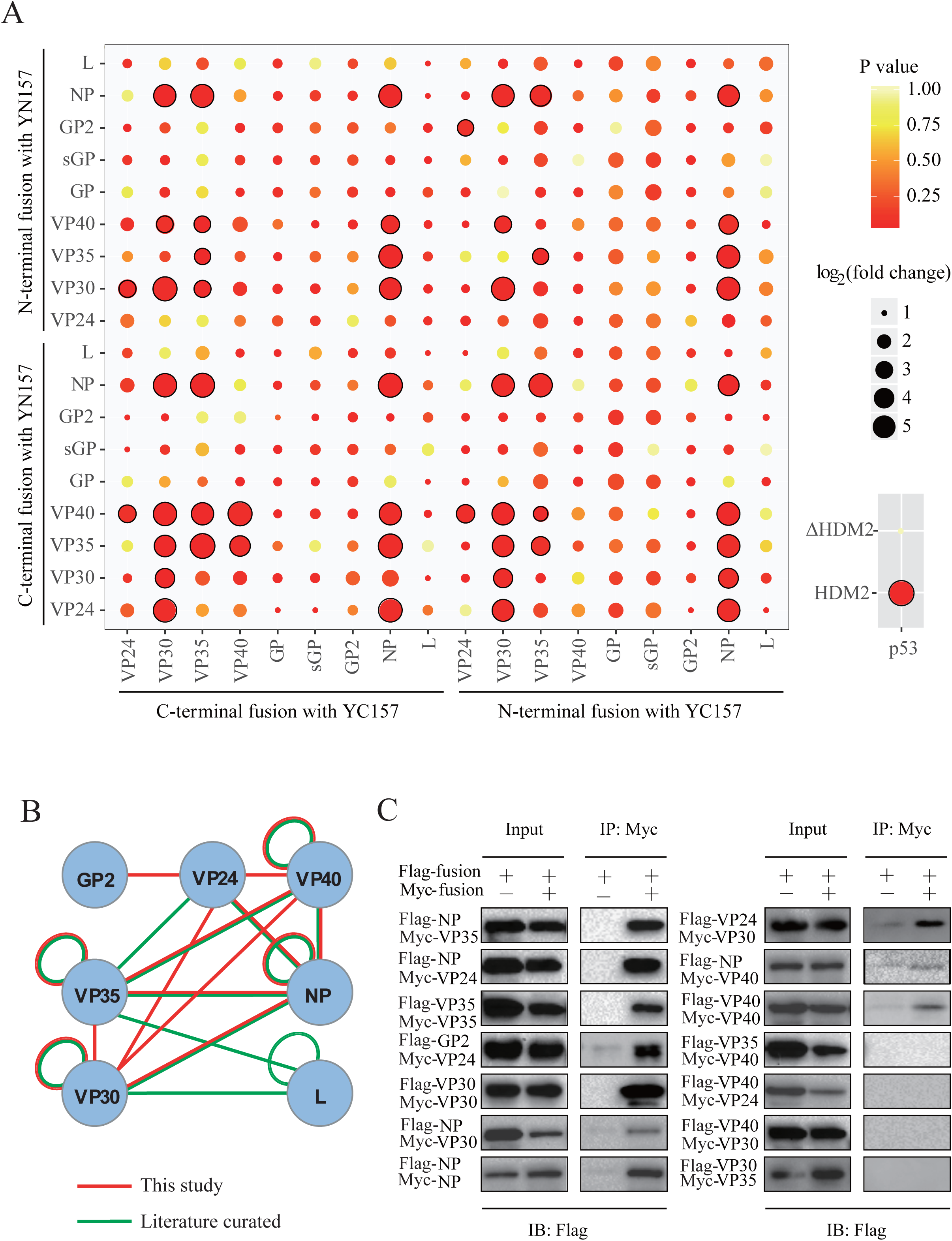
An intra-EBOV PPI network reveraled by BiFC. (A) An 18×18 cotransformation of yEGFP fragments tagged EBOV proteins were carried out as indicated, interactors are marked with a black circle, bJun and bFos/ΔbFos were used as control indicators. Data are presented as means±SD (n=3), Student’s t-test. (B) A global intra-PPI network of EBOV combined by our BiFC screening result with literature-curated datasets. (C) Validation of interactions among EBOV proteins by co-IP. EBOV proteins fused with Falg- or Myc-tags were expressed in HEK293 cells. Immunoprecipitations were performed using anti-Myc or anti-Flag antibodies.

### Screening p53 interactors of human library by using BiFC-seq

To further show that the BiFC-seq method is suitable for high-throughput screening, the well-known tumor suppressor p53 was used as a bait to screen its interactors against the human universal library fused with YN157 fragment (YN157-library). We used HDM2, a known partner of p53 as positive control [24]. The mutant form of HDM2 (ΔHDM2, with deletion of 1-139 aa), which lost its ability of binding to p53 was used as negative control. Fluorescent cells was effectively distinguished between positive and negative controls by yEGFP-BiFC (Figure S4A and B), we also tested various linkers [25] between p53 or HDM2 and yEGFP halves by yEGFP-BiFC method, we found that the (GGGGS)3 linker produced the highest fluorescence ratio among the four tested linkers by flow cytometry (Figure S4C and D). To screen p53 interactors against the library, the YN157-library was sequentially transformed into yeast competent cells containing p53 (YC157-p53 or p53-YC157) or their corresponding control groups with YC157 fragments (YC157-linker and linker-YC157), each transformation was performed in triplicate. After 72 hours of cultivation in SD liquid medium without tryptophan and leucine (SD-2), approximately 2,000 fluorescent cells out of 1×10^8^ cells were obtained for each p53 and control screening groups that were sorted by FACS (Figure S5), which is 10-100 times the number of positive colonies obtained with traditional Y2H screening.

To determine the identities of the positive colonies, PCR products of yeast colonies from each group were mixed and purified for NGS (Figure S6). The correlation coefficient of RNA-seq results from p53 groups among triplicate screenings was significantly higher than that of their corresponding controls (**Figure 3B**). After the different expression analysis by bioinformatics tools (**Figure 3A**), we identified 61 and 43 interactors for each of YC157-p53 and p53-YC157 screening groups. There are seven overlapped interactions between two groups (**Figure 3C** and Table S1). Totally 913 non-redundant interactions of p53 were recorded in BioGrid database [26]. To our knowledge this dataset is most comprehensive and we referred to these interactions as ‘known’ interactions. Twenty-one interactions of p53 were recaptured by this study. We estimated the sensitivity of BiFC-seq is ∼2% (21/913), which is similar to the large-scale Y2H screening (the sensitivity is ∼1%) [40]. The specificity of BiFC-seq was evaluated by co-IP assay. Totally 16 interactions of p53 were validated and 11 of them were re-confirmed. Thus the specificity of BiFC-seq is more than 60% (11/16) (**Figure 4C**). The sampling sensitivity was estimated by three repeat screens of YC157-p53 and p53-YC157. Totally 61 interactions were identified for the YC157-p53 group. Of these interactions, 45 interactions are commonly identified. More interactions were found with additional screens. The sampling sensitivity is 73.7% per screen. Similar result was obtained for the p53-YC157 data and the sampling sensitivity is 83.7%. Based on this data, we estimated at least six screens are needed to reach 90% saturation (Figure S7). In addition, the screening results were evaluated by PRINCESS tool, which was developed to assess the reliability of binary PPIs [27]. More than 64% of the interactions were of high confidence and had cutoff ratios greater than 2 (excluding the genes that were not recognizable) (Table S2). Then, we analyzed the function of p53 interactors by Gene Ontology, consistent with previous reports, the p53 partners participate in apoptosis, DNA damage response and translation *etc* (**Figure 4A** and Table S3). We further performed enrichment analysis of p53 by excluding 21 known p53 interactors. Most of the enriched GO terms are not changed, such as apoptosis, RNA catabolic process and translation *etc* (Figure S8 and Table S4). Four subnetworks of p53 interactors were enriched in the biological processes (**Figure 4B**), suggesting a mediator role of p53 among these subnetworks. The interaction of p53 with 5 interactors involved in the cilia formation subnetwork indicates that the p53 might be associated with dividing of centrosomes [28], which is consisted with previous reports in mouse cells [29], indicating the reliability of our screening results. To estimate the reproducibility of BiFC-seq, we analyzed the data of using p53 as bait. For the three replicates of YC157-p53, 45 of 61 interactions are consistently identified. For the p53-YC157 groups, 36 of 43 interactions are consistently identified in three replicates. These results indicate that for the library screening of BiFC-seq, about 77.9% are reproducible. A part of interactions of p53 were validated by co-IP and BiFC assay (Figure 4C and D).

**Figure 3.**
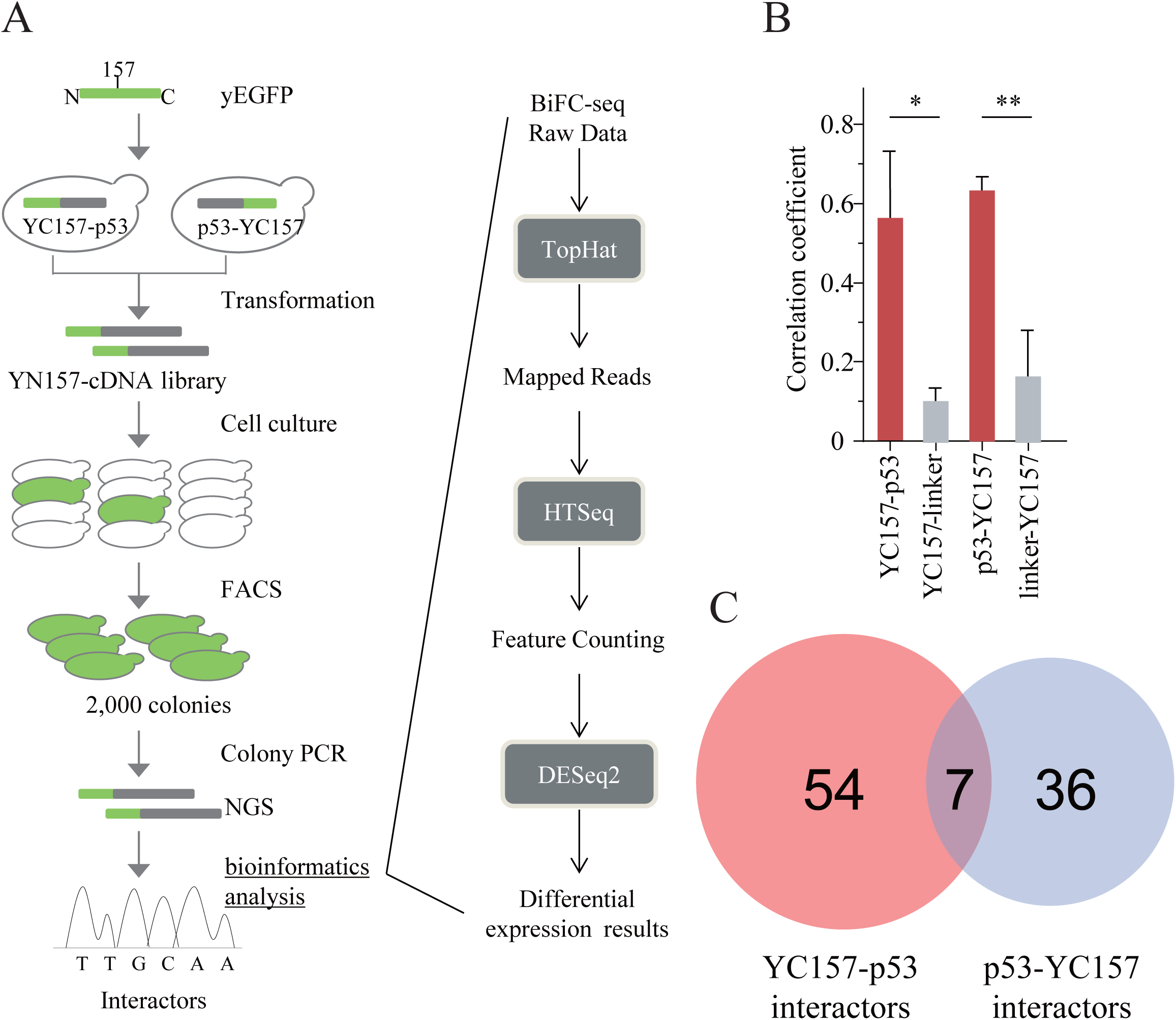
Identification of p53 interactors by BiFC-seq. (A) Flowchart of the BiFC-seq screening. The fluorescence-positive cells were used as templates to amplify the encoding genes of p53 binding partners. PCR products were sequenced by NGS and analyzed with bioinformatics tools to identify the interactors of p53. (B) Correlation analysis of p53 and their corresponding control screening groups. Data are presented as means±SD (n=3), \**P*<0.05, \*\**P*<0.01, Student’s t-test. (C) Number of screening interactors of p53 from YC157-p53 and p53-YC157 groups was presented.

**Figure 4.**
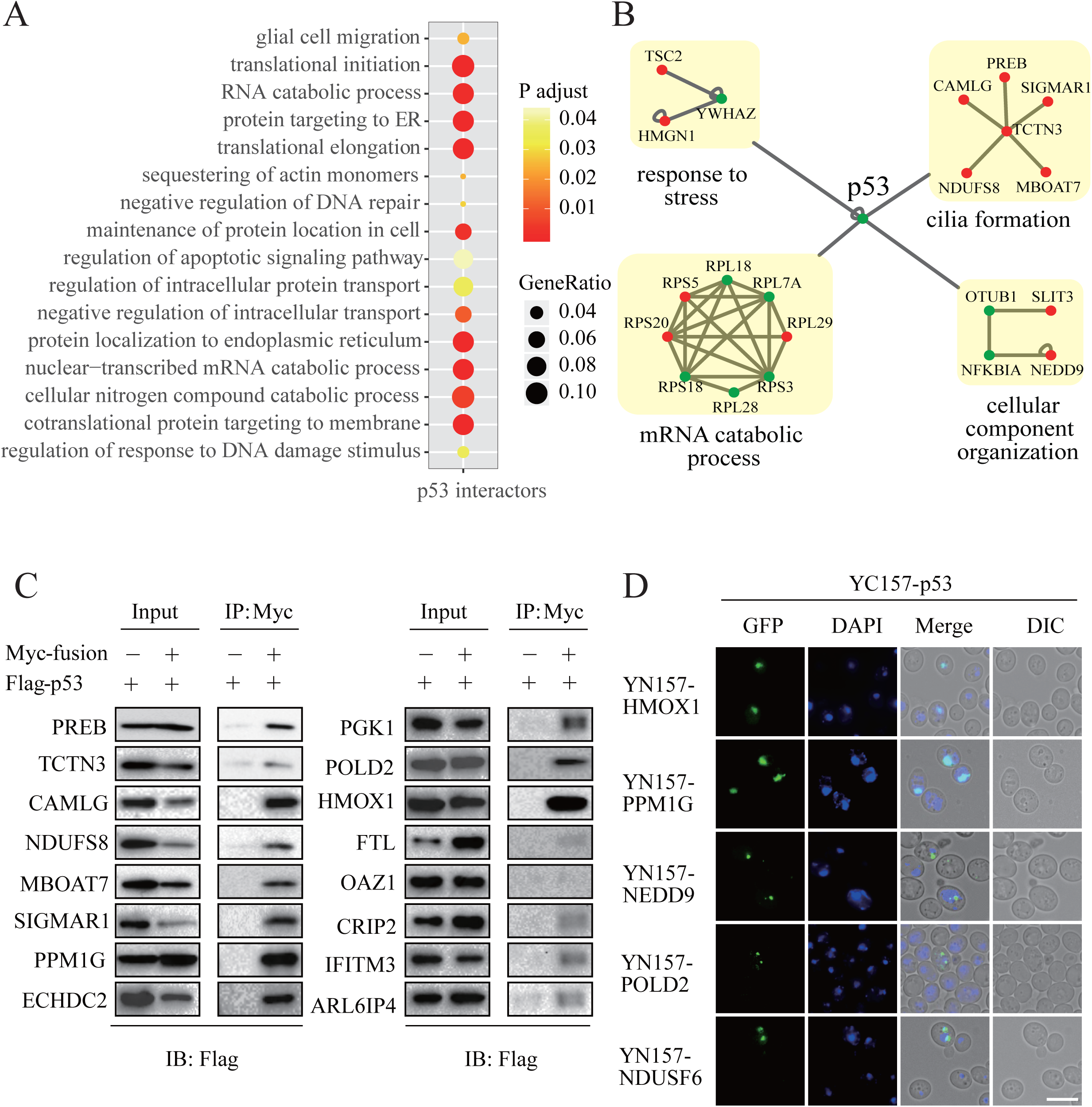
Analysis and validation of p53 interactors obtained by BiFC-Seq. (A) Selection of enriched biological processes for p53 screening results. (B) Subnetworks constituted by p53 interactors from the BiFC-seq results, blue and red dots represent interactors of p53 that reported and unreported from BioGrid database individually. (C) Validation of the p53 interactions by co-IP assays, wherein p53 was expressed as a Flag-tagged fusion and its partners were expressed as Myc-tagged fusions. HEK293 cells were transfected with the indicated constructs, immunoprecipitated and detected with anti-Flag antibody. (D) Validation of p53 interactions with novel screening partners by yEGFP-BiFC. p53 were cotransformed with interactors into yeast component cells as indicated, fluorescnece were detected under microscope 72 hours after cotransformation, scale bar: 10 μm.

The AP-MS method can be used to obtain protein complex information. Thus, we compared our data with a large-scale endogenous protein complex study [30]. A total of 50 high-confidence interactors of p53 were found by AP-MS, 13 of which were recorded in the BioGrid database (Table S5). Two of these interactions are overlapped with this study. AP-MS identifies members of stable complexes, whether they directly interact with the bait or not. In contrast to AP-MS method, BiFC-seq identifies binary interactions between pairs of proteins and tends to enrich transient interactions. Meanwhile, the stable binary interactions could be also identified by BiFC-seq method, such as interactions between ribosome proteins. The low overlap rate of the data from AP-MS and BiFC-seq might due to the differential features of technologies.

### A library to library interactome screening using BiFC-seq technology

Based on the aforementioned results, we extended the BiFC-seq screen to a genome-wide scale. Thus, we fused the human universal library to the C-terminal of YC157 fragment, yielding YC157-library pools. YN157-library and YC157-library pools were co-transformed into AH109 yeast component cells. After cultivation for 72 hours in SD-2 liquid medium followed by FACS analysis, fluorescent cells were divided into three groups according to their fluorescence intensities and approximately 3000 fluorescent cells were sorted out from each group (Figure S9). Yeast colonies were picked as PCR templates until their diameters reached 2-3 mm. To identify the interacting partners within the same fluorescent yeast cell from library screening by a single NGS experiment, we joined the encoding sequences of interacting pair into one PCR amplicon using stitching PCR strategy, which contains of two rounds of PCR (**Figure 5A**). In the first round, yeast colony PCR was conducted to amplify encoding sequences of interactors from YN157-library and YC157-library individually. In the second round, amplicons from the first round were used as templates for the stitching PCR to combine the amplicons together by their corresponding complementary upstream primers (Figure S10). To exclude most non-specific PCR amplicons from the first round PCR, stitching PCR products with more than 1000 bp fragments were mixed and purified for the NGS analysis using Illumina Miseq platform. Approximately 1 million ∼300 bp paired-end reads were generated from each group, the stitching amplicons that contain encoding sequences of interactors from both library pools linked by complementary sequences were extracted and mapped to the CDS regions of the human reference genome (GRCH38). Both of the gene encoding sequences from the same read that are in frame with the open reading frame (ORF) were considered to be interactors. After removing non-specific binding proteins that detected by screening interactors against YN157-library using YC157 fragments as baits, a network of 229 interactions among 205 proteins constituted by PPIs from three screening groups was generated and 12 interactions were reported in the BioGrid database (**Figure 6A** and Table S6). A minority of interactions were overlapped among three groups (**Figure 5B**), which indicated that a more wide range of PPI can be revealed by classifying the yeast cells with different fluorescent intensities. Totally, 137 interactions recognized by PRINCESS tool, 26 had high confidence with a cutoff ratio greater than 2 (Table S7). We classified these interactors by PANTHER [31]. Compared with all of proteins encoded in the human genome, these interactors have similar functional distribution, which suggest that the BiFC-seq method has no bias to obtain certain classes of PPI (**Figure 5C**). Furthermore, we found several clusters by superimposing PPI of this study onto BioGrid datasets (**Figure 6B-D**), such as gene expression, metabolic process, *etc*. For example, the FXYD domain containing ion transport regulator 2 (FXYD2), which was only found to be interacted with small molecule of cyclothiazide in BioGrid database [32], participates in protein metabolic process through interacting with translation related partners (**Figure 6E**). The resources of protein-protein interactions are very useful for therapeutic discovery, such as drug repurposing. The potential drugs are identified by network-based methodologies. For example, dozens of candidate drugs were prioritized for SARS-CoV-2 by using the interactome of virus and host [33],[34]. For our interesting, the connections of ANXA2-DAXX and SNRPB2-PSMB6 were involved in the viral infection processes and regulation of immune process (**Figure 6F and G**). These results demonstrate the great potential of BiFC-seq technology in genome-wide interactome screening.

**Figure 5.**
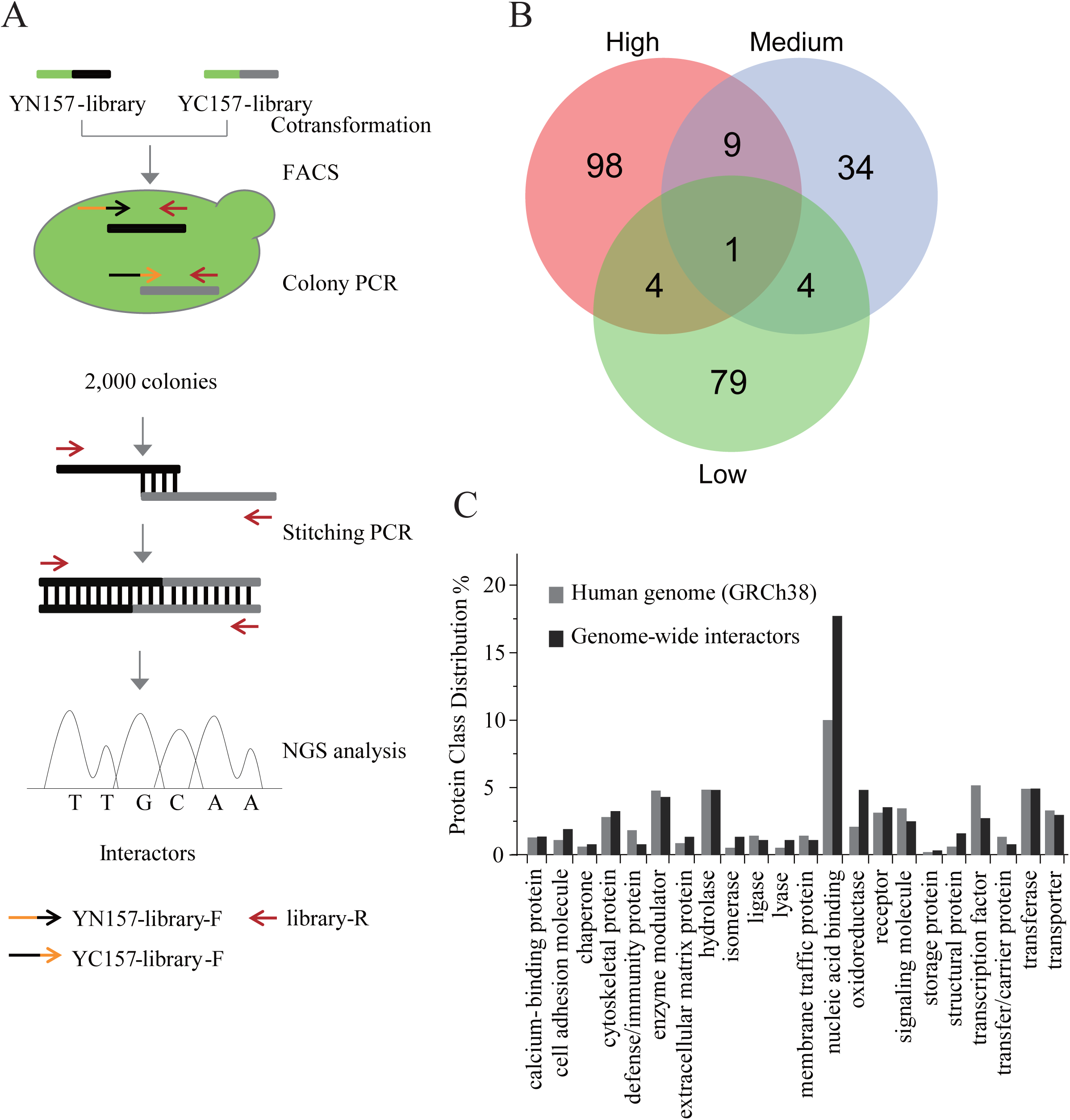
Library to library screening by BiFC-seq. (A) Experimental workflow of genome-wide interactome screening. (B) Venn diagram of interaction overlaps among three screening groups. (C) Protein class distribution of genome-wide interactors along with that of human genome.

**Figure 6.**
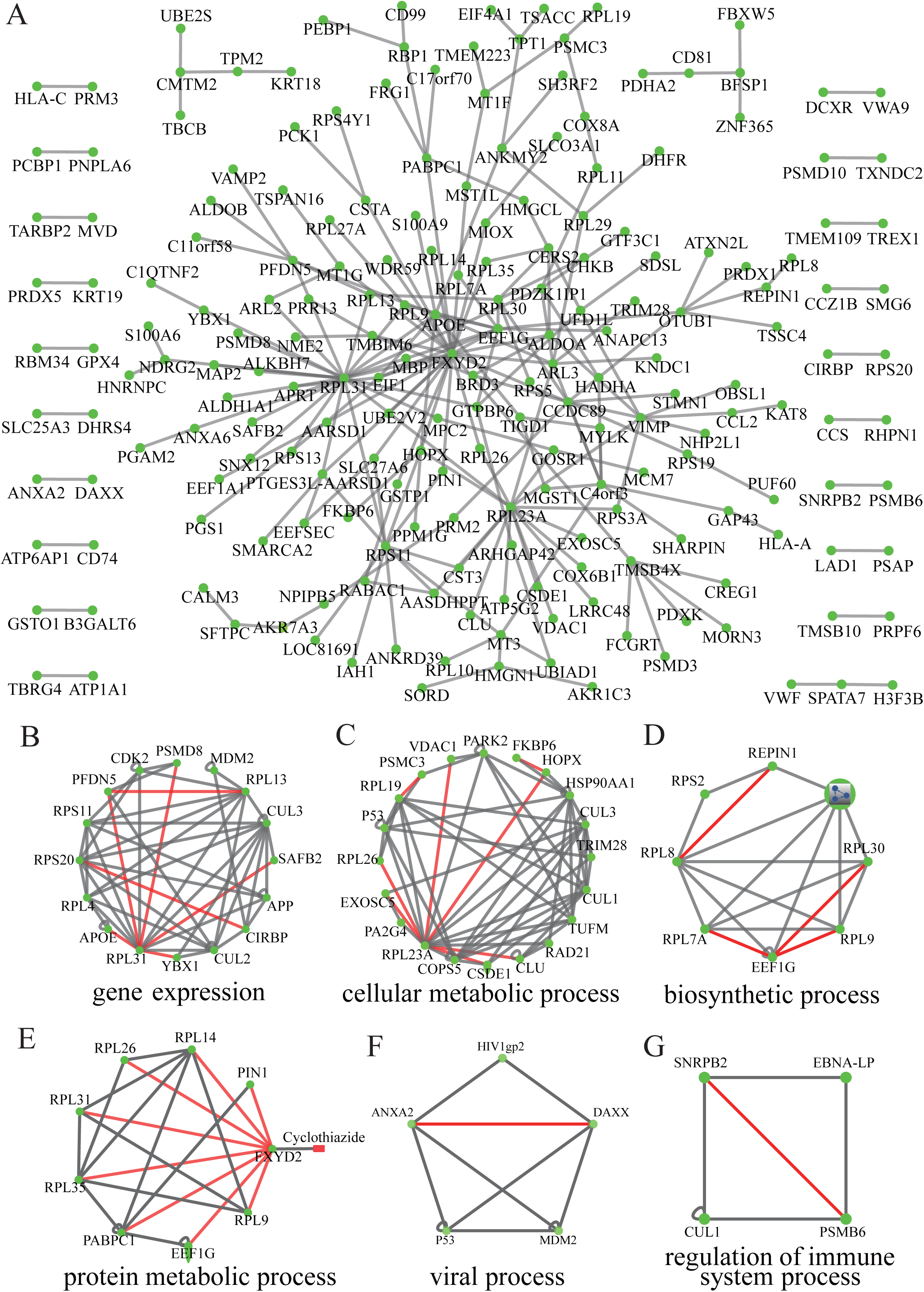
A PPI networks based on BiFC-seq. (A) A global network constituted by library to library screening results by BiFC-seq. (B-G) Various bioligical processes are presented by intergrating BiFC-seq interactions with BioGrid dataset, grey and red lines represent interactions from BioGrid database and BiFC-seq results individually.

## Conclusion

To our knowledge, the large-scale PPI studies using BiFC method were mainly based on optimized Venus protein, which is a well-worked system in mammalian cells. However, when applied in other organisms, such as yeast and plant, it has low signal-to-noise ratio and high background signal. Here, we developed a yEGFP-based yeast BiFC method with a higher sensitivity and specificity. Compared with traditional Y2H method, the baits are neither required to import into nucleus to activate reporters nor needed to test self-activation before screening, which has extended the scope of PPI screenings.

To validate the feasibility of using yEGFP-BiFC method in interactome research, we searched the interactors of the nine encoding proteins of EBOV. The results revealed a network of 14 interactions among 6 proteins with a high coverage of reported interactions (9/14) and 10 interactions were validated by co-IP assays. By integrating the PPIs of EBOV in this study with reported interactions, we generated a global EBOV PPI network, which provides new insights into pathogenesis from the perspective of the intraviral network. Compared with genetics approaches, such as gene-trap insertional mutagenesis [35], RNAi [36] or CRISPR-CAS9 screening [37], which reveals gene-gene interactions and most of them are genetics interactions, the protein-protein interactions identified by BiFC-seq are mostly physical and binary interactions. The interactions from these two kinds of methods are complementary.

Next, we justified the feasibility of BiFC-seq in a high-throughput PPI screenings, p53 was employed to screen against a human cDNA-library. We obtained 21 reported and 76 novel interactors of p53 from 97 partners. Seven interactors were overlapped for the N-terminal and C-terminal labeling p53, which demonstrated the necessity of using different fusion constructs in PPI screening. For our interesting, we found four enriched subnetworks formed by p53 interactors. The involvement of p53 in the cilia formation suggests its role in restraining overgrowth of centrosomes, which is a prerequisite for cancer development in mice [38]. We also confirmed 11 interactions of p53 by co-IP assay, which suggests the high-confidence of these interactions. Moreover, we found some membrane proteins, such as calcium modulating ligand (CAMLG), secretory carrier membrane protein 3 (SCAMP3) and interferon induced transmembrane protein 3 (IFITM3) that interact with p53, which have low possibility to be detected by traditional Y2H method.

Compared with the widely used Y2H and AP-MS technologies for genome-wide interactome mapping, the BiFC-seq approach is highly sensitive, has excellent screening capability, and incurs relatively low financial and labor costs. We obtained a total of 97 p53 interactions, of which 21 were previously known, whereas only 51 interactions were obtained and recorded in the BioGrid database for three proteomic-scale Y2H screenings using p53 as bait (Table S8) [39]-[41]. Thus, the BiFC-seq approach had a higher sampling sensitivity than the traditional Y2H library or array screening approaches.

Furthermore, we extended the BiFC-seq method to a genome-wide interactome screening by pooling YN157 and YC157 tagged human cDNA libraries together. By analyzing 9000 fluorescent cells from three screening groups with different fluorescence intensities, we obtained 229 interactions of 205 proteins in a single BiFC-seq screening. The interactions participate in various biological processes, such as gene expression, metabolism and viral infection, which suggests that the BiFC-seq technology is high-sensitive to delineate genome-wide interactome in future application. We have compared the interactions identified by BiFC-seq with non-interacting protein pairs from Negatome dataset [42]. The transcriptomics data of 74 tissues were from human protein atlas database (HPA) [43]. The interactions with absolute value of spearman correlation coefficient (SCC) greater than 0.5 were considered positive co-expression pairs. Compared with Negatome data, the p53 interactors were more likely to be co-expressed. The ratio of co-expressed interactions of p53 is 20.8%, while for Negatome data the value is 11.4%. For the interactions of library screening, the ratio (13.1%) is slightly greater than that of Negatome data (Table S9).

Compare with traditional approaches, such as Y2H and AP-MS technologies, the BiFC-seq has its limitations. Most of the Y2H method employs more than one reporters while BiFC-seq depends on EGFP reporter only, which might lead to more false positives. Moreover, in contrast to AP-MS, BiFC-seq is amenable in yeast cells, which results in the interactions that depends on post-translational modification or mediated by other proteins couldn’t be detected. AP-MS is performed *in vivo*, which investigates protein complexes under physiological conditions.

Recently, the CrY2H-seq technology was introduced, which uses a Cre recombinase reporter and next-generation DNA sequencing to identify interactions [44]. CrY2H-seq is a highly scalable screening approach to detect binary protein–protein interactions in yeast. Similar to BiFC-seq, CrY2H-seq is cost-effective and time-efficient, which greatly increases interaction detection rates. On the other hand, since CrY2H-seq is based on yeast transcription factor that the self-activation of considerable baits is inevitable (∼16% baits are self-activated). CrY2H-seq uses auxotrophic rescue reporter, which usually takes more time (2∼5 day) to allow the growth of yeast. BiFC-seq uses a fluorescent reporter, and the signal can be detected within 2 days in yeast. Compare with CrY2H-seq, a single reporter of BiFC-seq might increase the false positives.

It should be noted that only limited interactions were identified for the human interactome by a library-library screening in this study. The major reasons are as following. First, the quality of cDNA library is limited. The high abundance cDNA, such as ribosomal genes, accounts for the top list the identified interactions. The variation of cDNA isoforms and truncations also restrict the number of unique interactions after removing the redundant interactions. Second, it remains difficult to clone all of the human genes to BiFC-seq vectors, which might greatly reduce the redundant interactions and false positives from truncations or out-of-frame expressions. They cloned 1956 genes to CrY2H-seq vectors and 8577 interactions were obtained, which partially reflects the importance of cloning the full-length genes. Third, we sorted 9000 positive colonies, which represent about 200 interactions. To obtain more interactions by library to library screening, more colonies need to be identified in future.

In summary, we have developed a BiFC-seq method, which is amenable in high-throughput PPI screening by using FACS. The sensitivity of BiFC-seq was substantially increased by coupling with NGS. This technology is highly sensitive for screening physical binary PPIs and can be easily applied to other large-scale interactome mapping studies.

## Authors’ contributions

L.S. performed BiFC screens. Y.Z. performed the co-IP assays. Y. L. performed the PCR assays. L.S., C.J., Y.Y., C.T., D.L., L.Z., and J.W. contributed to the data analysis. M.N. and X.B. performed the NGS experiment. L.S. and J.W. wrote the manuscript. F.H. and J.W. conceived and supervised the study.

## Acknowledgments

We thank Professor Alistair from University of Aberdeen to kindly provide pUC19-YGFP plasmid. This work was supported by grants from the the National Key Research and Development Program (No. 2017YFA0505700) and the grant from the National Key Lab of Proteomics (SKLP-K201805, SKLP-K201804).

## Supplementary material

**Figure S 1**

**Expression of yEGFP in yeast cells**.

Intact yEGFP or YN157/YC157 fragments were transformed into yeast cells that were then cultured at 30°C for 24 hours in SD-2 medium and incubated at 4°C for approximately 48 hours for fluorophore maturation before observation, scale bar: 10 μm. WL: whole length.

**Figure S 2.**

**Expression of EBOV proteins fused with fluorescent fragments**.

N-terminal flag tagged EBOV proteins fused with fluorescent fragments as indicated, expression of fused protein are detected by anti-flag antibody.

**Figure S 3.**

**Validation of intra EBOV PPIs by yEGFP-BiFC**.

To validate the intra EBOV PPIs detected by flowcytometry, EBOV proteins with yEGFP fragment tags were contransfomed into yeast cell, the fluorescence of cells were observed under fluorescent microscope

**Figure S 4.**

**Validation of the interaction of HDM2 and p53 by BiFC**.

(A) p53 and HDM2/ΔHDM2 were cotransformed into yeast cells as indicated, the fluorescent yeast cells were observed under fluorescnece microscope, scale bar: 10 μm (B) fluorescent cells from (a) were analyzed by flow cytometry. (C,D) Various peptides that used to link YN157-HDM2 and YC157-p53 as indicated were check for their fluorescent cell ratio 72 hours after cotransformation by flow cytometry. Data are presented as means ± SD (n=3). Error bars, standard deviation. **\***, P<0.05, n.s., non significant.

**Figure S 5**

**The fluorescent yeast cells from p53 screening were sorted by FACS**. YN157-tagged human universal library was sequentially transformed into yeast cells containing p53 (YC157-p53 and p53-YC157) and control (YC157-linker and liker-YC157) groups. The fluorescent cells were sorted 72 hours after transformation. Approximately 2000 fluorescent cells were sorted from the P2 gate of each group.

**Figure S 6**

**Yeast colony PCR.**

**(A)** Fluorescent yeast colonies were used as templates to amplify the cDNA inserts. A portion of the PCR products were used for electrophoresis as indicated. **(B)** The electrophoresis results of the purified PCR products for each group. Mr: DNA2000 marker.

**Figure S 7**

**Number of interactions detected after each screen using p53 as bait by BiFC-seq**

**Figure S 8**

**The enriched Gene Ontology terms (biological processes) for unreported interactors of p53**.

**Figure S 9**

**The fluorescent yeast cells from library to library screening were sorted by FACS**.

The fluorescent yeast cells were sorted out into three groups according to their fluorescence intensities.

**Figure S 10**

**Electrophoresis of stitching PCR products**.

Gel electrophoresis of stitching PCR products. N-library and C-library represent yeast colony PCR products derivated from YN157-library and YC157-library controls. N-library +C-library represents stitching PCR products from test groups. Mr: DNA15000 marker.

**Table S 1.**

**The screening results of using p53 as bait by BiFC-seq.**

**Gene Symbol**: NCBI ENTREZ Symbols of the interactors.

**Group**: Specific highly expressed transcripts in each group according of p53-YC157 and YC157-p53. p53 indicates the overlap interactors using p53-YC157 and YC157-p53 as baits.

**Table S 2.**

**Evaluation of the reliability of the p53 interactions by PRINCESS**.

Confidence scores were measured by a Bayesian approach that combines biological evidence from multiple sources. #interaction: the number of interactions in each category; LR: likelihood ratio.

**Table S 3.**

**The Gene Ontology analysis of p53 interactors.**

**GO ID**: Gene Ontology ID.

**Description**: description of the GO term.

**GeneRatio**: ratio of the number of genes in a GO category to all the genes in the sample.

**BgRatio**: ratio of the number of genes in a GO category to the background genes.

**Gene ID**: NCBI ENTREZ GeneID of the genes belonging to a GO category.

**Count**: number of all genes belonging to a GO category.

**Table S 4.**

**The Gene Ontology analysis of unreported interactors of p53.**

**GO ID**: Gene Ontology ID.

**Description**: description of the GO term.

**GeneRatio**: ratio of the number of genes in a GO category to all the genes in the sample.

**BgRatio**: ratio of the number of genes in a GO category to the background genes.

**Gene ID**: NCBI ENTREZ GeneID of the genes belonging to a GO category.

**Table S 5.**

**The reported interactions in three proteomic-scale Y2H screenings using p53 as bait**.

Official Symbol_a/Official Symbol_b: NCBI ENTREZ Symbol of a or b gene; Gene ID_a/Gene ID_b: NCBI ENTREZ GeneID of a or b gene.

**Table S 6.**

**A genome-wide BiFC-seq PPIs screening results**.

The reads of the same amplicon derived from YN157-library and YC157-library were mapped to the genome CDS regions individually. The gene symbols of the interactors from three screening groups were shown.

**Table S 7.**

**Evaluation of the reliability of the genome-wide PPI screening results by PRINCESS**.

Confidence scores were measured by a Bayesian approach that combines biological evidence from multiple sources. #interaction: the number of interactions in each category; LR: likelihood ratio

**Table S 8.**

**The p53 interactors in a large-scale AP-MS screening using p53 as bait**.

Official Symbol_a/Official Symbol_b: NCBI ENTREZ Symbol of a or b gene Gene ID_a/Gene ID_b: NCBI ENTREZ GeneID of a or b gene

**Table S 9.**

Co-expression analysis of PPI pairs identified by BiFC-seq. BiFC-seq identified PPI pairs were compared with human negatome dataset, which is a collection of protein and domain pairs that are unlikely engaged in direct physical interactions. Those with absolute value of spearman correlation coefficient (SCC) greater than 0.5 were considered co-expression pairs.

## References

[1] Huttlin EL, Bruckner RJ, Paulo JA, Cannon JR, Ting L, Baltier K, et al. Architecture of the human interactome defines protein communities and disease networks. Nature 2017;545:505–9.

[2] Shah PS, Link N, Jang GM, Sharp PP, Zhu T, Swaney DL, et al. Comparative Flavivirus-Host Protein Interaction Mapping Reveals Mechanisms of Dengue and Zika Virus Pathogenesis. Cell 2018;175:1931–45 e18.

[3] Choi SG, Richardson A, Lambourne L, Hill DE, Vidal M. Protein Interactomics by Two-Hybrid Methods. Methods Mol Biol 2018;1794:1–14.

[4] Vidal M, Fields S. The yeast two-hybrid assay: still finding connections after 25 years. Nat Methods 2014;11:1203–6.

[5] Menche J, Sharma A, Kitsak M, Ghiassian SD, Vidal M, Loscalzo J, et al. Disease networks. Uncovering disease-disease relationships through the incomplete interactome. Science 2015;347:1257601.

[6] Venkatesan K, Rual JF, Vazquez A, Stelzl U, Lemmens I, Hirozane-Kishikawa T, et al. An empirical framework for binary interactome mapping. Nat Methods 2009;6:83–90.

[7] Sung MK, Lim G, Yi DG, Chang YJ, Yang EB, Lee K, et al. Genome-wide bimolecular fluorescence complementation analysis of SUMO interactome in yeast. Genome Res 2013;23:736–46.

[8] Miller KE, Kim Y, Huh WK, Park HO. Bimolecular Fluorescence Complementation (BiFC) Analysis: Advances and Recent Applications for Genome-Wide Interaction Studies. J Mol Biol 2015;427:2039–55.

[9] Lee OH, Kim H, He Q, Baek HJ, Yang D, Chen LY, et al. Genome-wide YFP fluorescence complementation screen identifies new regulators for telomere signaling in human cells. Mol Cell Proteomics 2011;10:M110 001628.

[10] Trapnell C, Roberts A, Goff L, Pertea G, Kim D, Kelley DR, et al. Differential gene and transcript expression analysis of RNA-seq experiments with TopHat and Cufflinks. Nat Protoc 2012;7:562–78.

[11] Anders S, Pyl PT, Huber W. HTSeq--a Python framework to work with high-throughput sequencing data. Bioinformatics 2015;31:166–9.

[12] Love MI, Huber W, Anders S. Moderated estimation of fold change and dispersion for RNA-seq data with DESeq2. Genome Biol 2014;15:550.

[13] Cormack BP, Bertram G, Egerton M, Gow NA, Falkow S, Brown AJ. Yeast-enhanced green fluorescent protein (yEGFP): a reporter of gene expression in Candida albicans. Microbiology 1997;143 (Pt 2):303–11.

[14] Gustin JK, Bai Y, Moses AV, Douglas JL. Ebola Virus Glycoprotein Promotes Enhanced Viral Egress by Preventing Ebola VP40 From Associating With the Host Restriction Factor BST2/Tetherin. J Infect Dis 2015;212 Suppl 2:S181–90.

[15] Garcia-Dorival I, Wu W, Dowall S, Armstrong S, Touzelet O, Wastling J, et al. Elucidation of the Ebola virus VP24 cellular interactome and disruption of virus biology through targeted inhibition of host-cell protein function. J Proteome Res 2014;13:5120–35.

[16] Baseler L, Chertow DS, Johnson KM, Feldmann H, Morens DM. The Pathogenesis of Ebola Virus Disease. Annu Rev Pathol 2017;12:387–418.

[17] Groseth A, Charton JE, Sauerborn M, Feldmann F, Jones SM, Hoenen T, et al. The Ebola virus ribonucleoprotein complex: a novel VP30-L interaction identified. Virus Res 2009;140:8–14.

[18] Huang Y, Xu L, Sun Y, Nabel GJ. The assembly of Ebola virus nucleocapsid requires virion-associated proteins 35 and 24 and posttranslational modification of nucleoprotein. Mol Cell 2002;10:307–16.

[19] Reid SP, Cardenas WB, Basler CF. Homo-oligomerization facilitates the interferon-antagonist activity of the ebolavirus VP35 protein. Virology 2005;341:179–89.

[20] Watanabe S, Noda T, Kawaoka Y. Functional mapping of the nucleoprotein of Ebola virus. J Virol 2006;80:3743–51.

[21] Hartlieb B, Muziol T, Weissenhorn W, Becker S. Crystal structure of the C-terminal domain of Ebola virus VP30 reveals a role in transcription and nucleocapsid association. Proc Natl Acad Sci U S A 2007;104:624–9.

[22] Liu Y, Stone S, Harty RN. Characterization of filovirus protein-protein interactions in mammalian cells using bimolecular complementation. J Infect Dis 2011;204 Suppl 3:S817–24.

[23] Johnson RF, McCarthy SE, Godlewski PJ, Harty RN. Ebola virus VP35-VP40 interaction is sufficient for packaging 3E-5E minigenome RNA into virus-like particles. J Virol 2006;80:5135–44.

[24] Kruse JP, Gu W. Modes of p53 regulation. Cell 2009;137:609–22.

[25] Petschnigg J, Groisman B, Kotlyar M, Taipale M, Zheng Y, Kurat CF, et al. The mammalian-membrane two-hybrid assay (MaMTH) for probing membrane-protein interactions in human cells. Nat Methods 2014;11:585–92.

[26] Chatr-Aryamontri A, Breitkreutz BJ, Oughtred R, Boucher L, Heinicke S, Chen D, et al. The BioGRID interaction database: 2015 update. Nucleic Acids Res 2015;43:D470–8.

[27] Li D, Liu W, Liu Z, Wang J, Liu Q, Zhu Y, et al. PRINCESS, a protein interaction confidence evaluation system with multiple data sources. Mol Cell Proteomics 2008;7:1043–52.

[28] Gupta GD, Coyaud E, Goncalves J, Mojarad BA, Liu Y, Wu Q, et al. A Dynamic Protein Interaction Landscape of the Human Centrosome-Cilium Interface. Cell 2015;163:1484–99.

[29] Bazzi H, Anderson KV. Acentriolar mitosis activates a p53-dependent apoptosis pathway in the mouse embryo. Proc Natl Acad Sci U S A 2014;111:E1491–500.

[30] Malovannaya A, Lanz RB, Jung SY, Bulynko Y, Le NT, Chan DW, et al. Analysis of the human endogenous coregulator complexome. Cell 2011;145:787–99.

[31] Mi H, Muruganujan A, Thomas PD. PANTHER in 2013: modeling the evolution of gene function, and other gene attributes, in the context of phylogenetic trees. Nucleic Acids Res 2013;41:D377–86.

[32] Imming P, Sinning C, Meyer A. Drugs, their targets and the nature and number of drug targets. Nat Rev Drug Discov 2006;5:821–34.

[33] Zhou Y, Hou Y, Shen J, Huang Y, Martin W, Cheng F. Network-based drug repurposing for novel coronavirus 2019-nCoV/SARS-CoV-2. Cell Discov 2020;6:14.

[34] Gordon DE, Jang GM, Bouhaddou M, Xu J, Obernier K, White KM, et al. A SARS-CoV-2 protein interaction map reveals targets for drug repurposing. Nature 2020.

[35] Cheng F, Murray JL, Zhao J, Sheng J, Zhao Z, Rubin DH. Systems Biology-Based Investigation of Cellular Antiviral Drug Targets Identified by Gene-Trap Insertional Mutagenesis. PLoS Comput Biol 2016;12:e1005074.

[36] Laufer C, Fischer B, Billmann M, Huber W, Boutros M. Mapping genetic interactions in human cancer cells with RNAi and multiparametric phenotyping. Nat Methods 2013;10:427–31.

[37] Shen JP, Zhao D, Sasik R, Luebeck J, Birmingham A, Bojorquez-Gomez A, et al. Combinatorial CRISPR-Cas9 screens for de novo mapping of genetic interactions. Nat Methods 2017;14:573–6.

[38] Coelho PA, Bury L, Shahbazi MN, Liakath-Ali K, Tate PH, Wormald S, et al. Over-expression of Plk4 induces centrosome amplification, loss of primary cilia and associated tissue hyperplasia in the mouse. Open Biol 2015;5:150209.

[39] Wang J, Huo K, Ma L, Tang L, Li D, Huang X, et al. Toward an understanding of the protein interaction network of the human liver. Mol Syst Biol 2011;7:536.

[40] Rual JF, Venkatesan K, Hao T, Hirozane-Kishikawa T, Dricot A, Li N, et al. Towards a proteome-scale map of the human protein-protein interaction network. Nature 2005;437:1173–8.

[41] Stelzl U, Worm U, Lalowski M, Haenig C, Brembeck FH, Goehler H, et al. A human protein-protein interaction network: a resource for annotating the proteome. Cell 2005;122:957–68.

[42] Smialowski P, Pagel P, Wong P, Brauner B, Dunger I, Fobo G, et al. The Negatome database: a reference set of non-interacting protein pairs. Nucleic Acids Res 2010; 38:D540–4.

[43] Uhlen M, Fagerberg L, Hallstrom BM, Lindskog C, Oksvold P, Mardinoglu A, et al. Proteomics. Tissue-based map of the human proteome. Science 2015; 347:1260419.

[44] Trigg SA, Garza RM, MacWilliams A, Nery JR, Bartlett A, Castanon R, et al. CrY2H-seq: a massively multiplexed assay for deep-coverage interactome mapping. Nat Methods 2017; 14:819–25.

