## Supplementary figures and images for "A yeast BiFC-seq method for genome-wide interactome mapping"

### supplementary figure 1

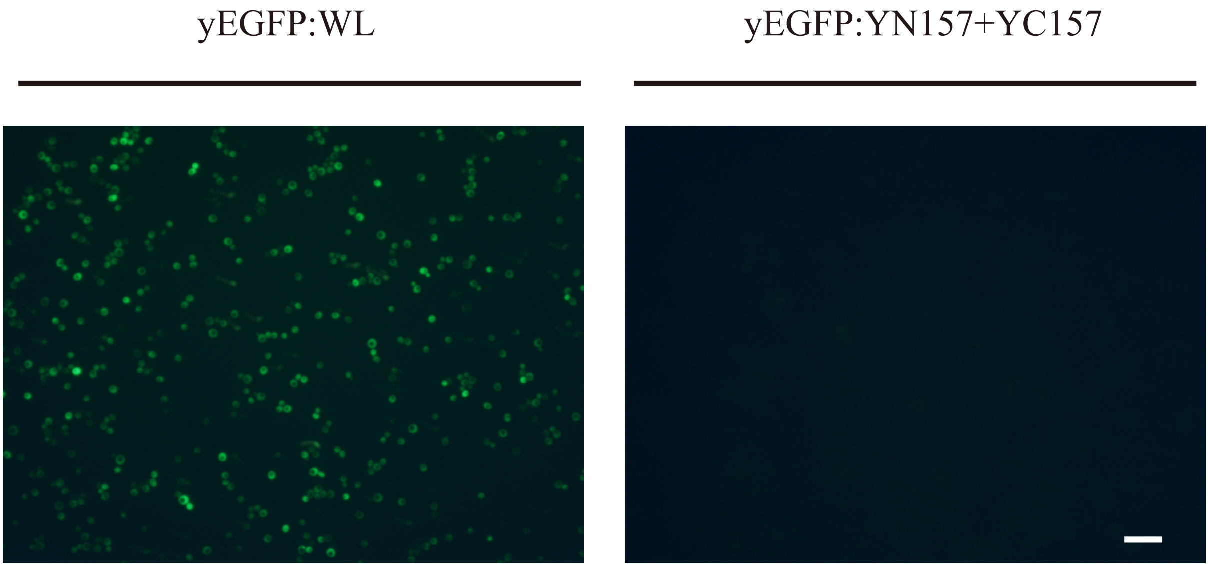


**Supplementary Figure 1**

### supplementary figure 2

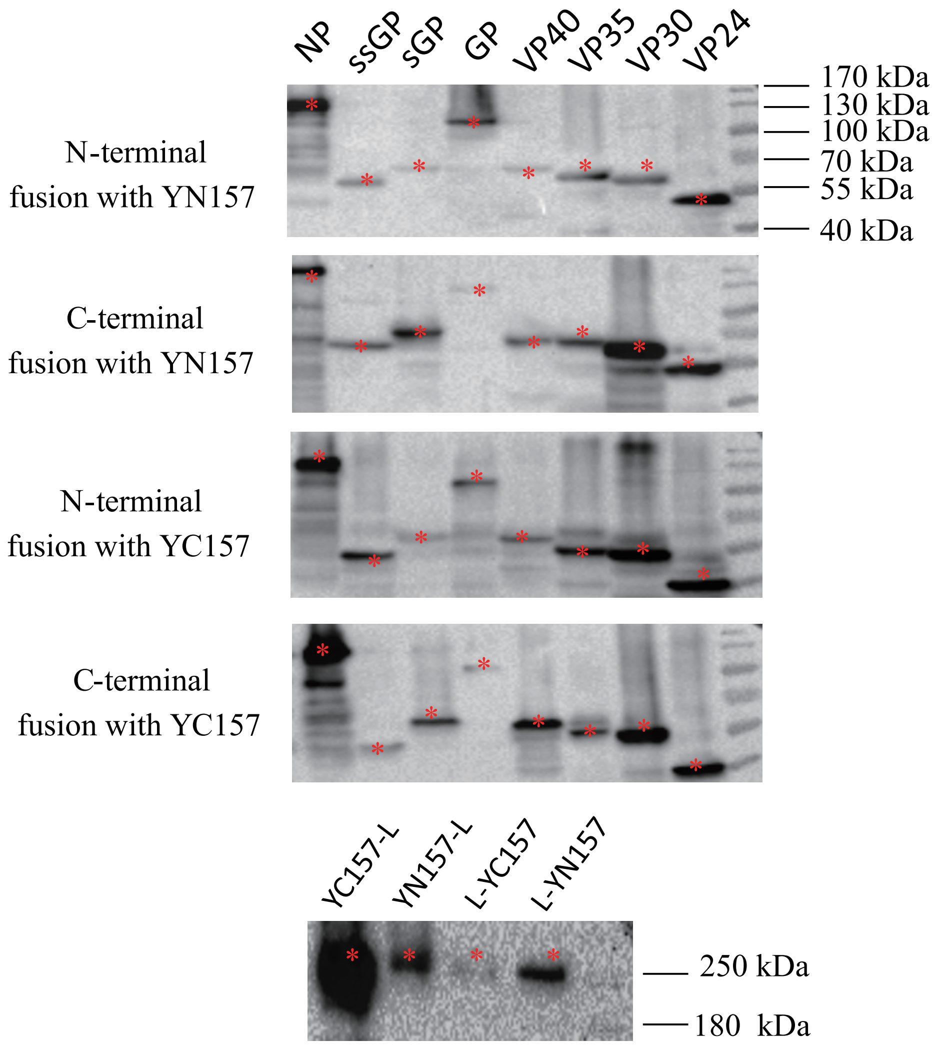


**Supplementary Figure 2**

### supplementary figure 4

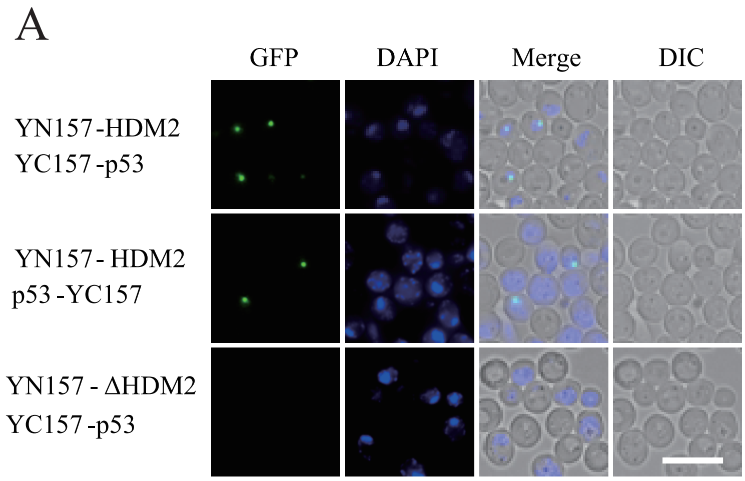


C

| | |
|---|---|
| Linker1 | (GGGGGS)3 |
| Linker2 | (GGGGGS)2 |
| Linker3 | RSIAT |
| Linker4 | RPACKIPNDLQKVMN |

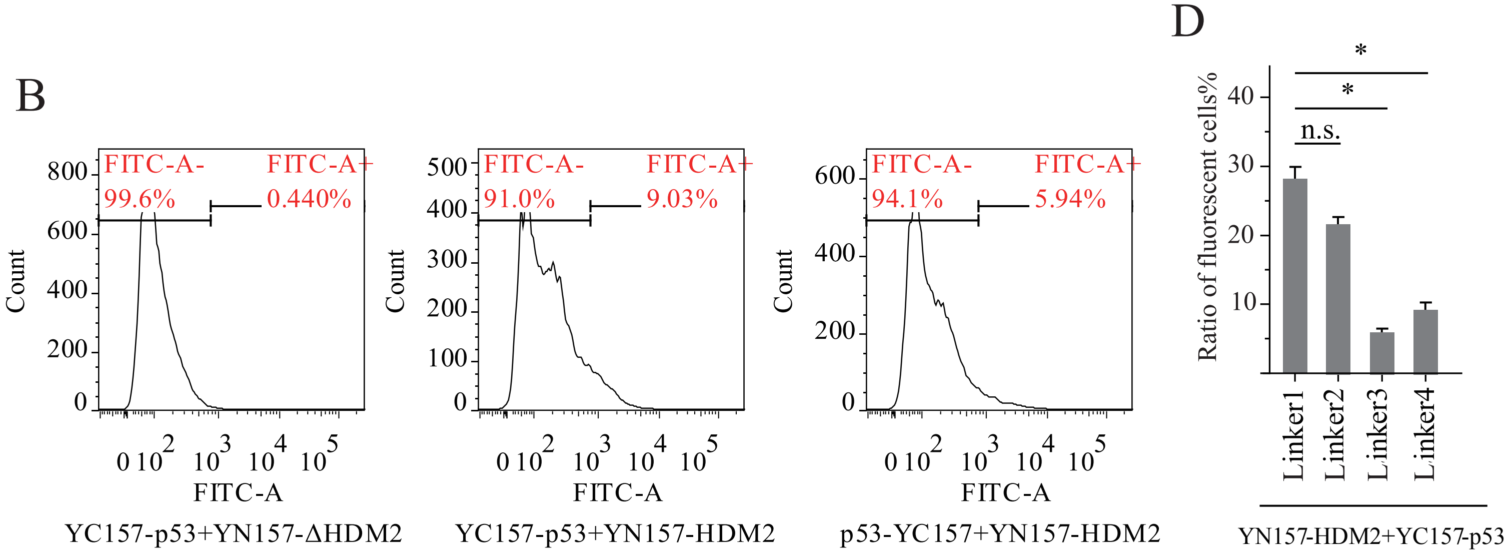


**Supplementary Figure 4**

### supplementary figure 5

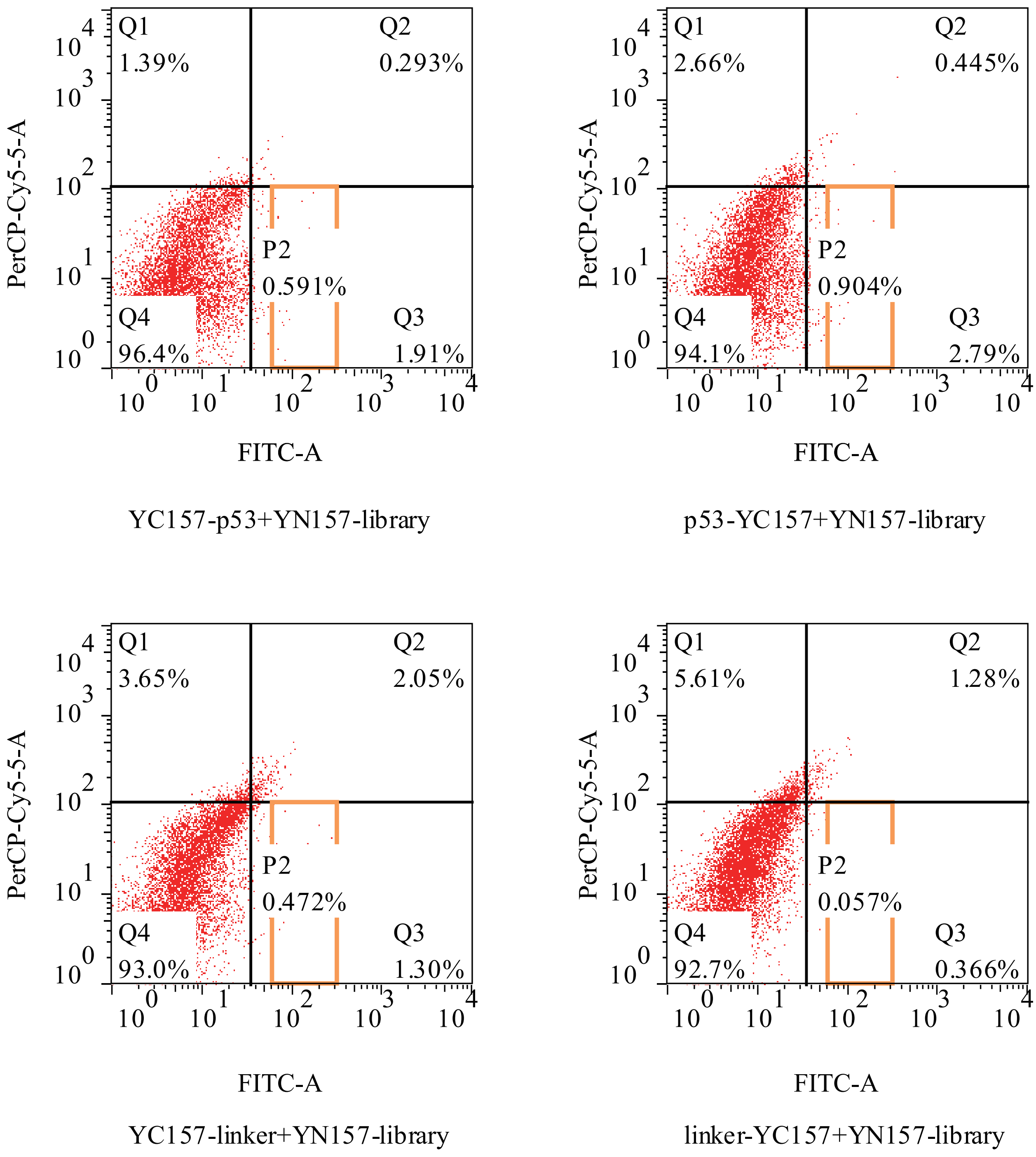


**Supplementary Figure 5**

### supplementary figure 6

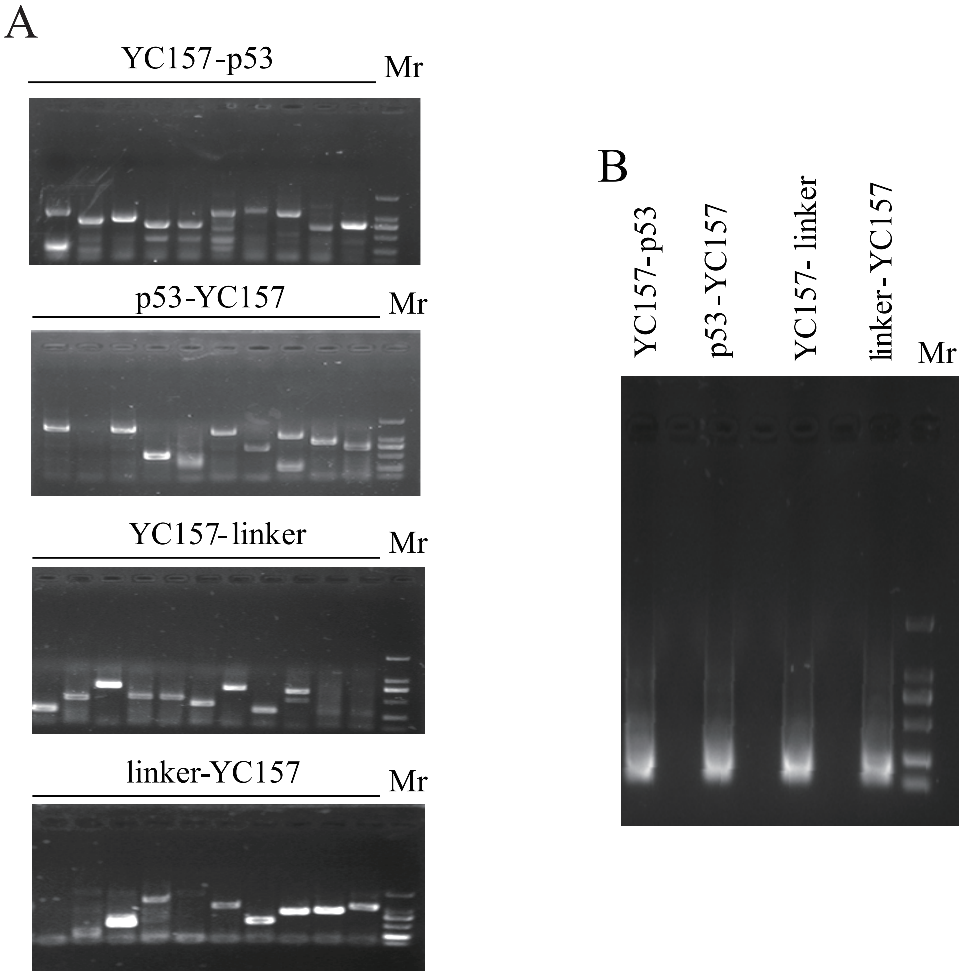


**Supplementary Figure 6**

### supplementary figure 7

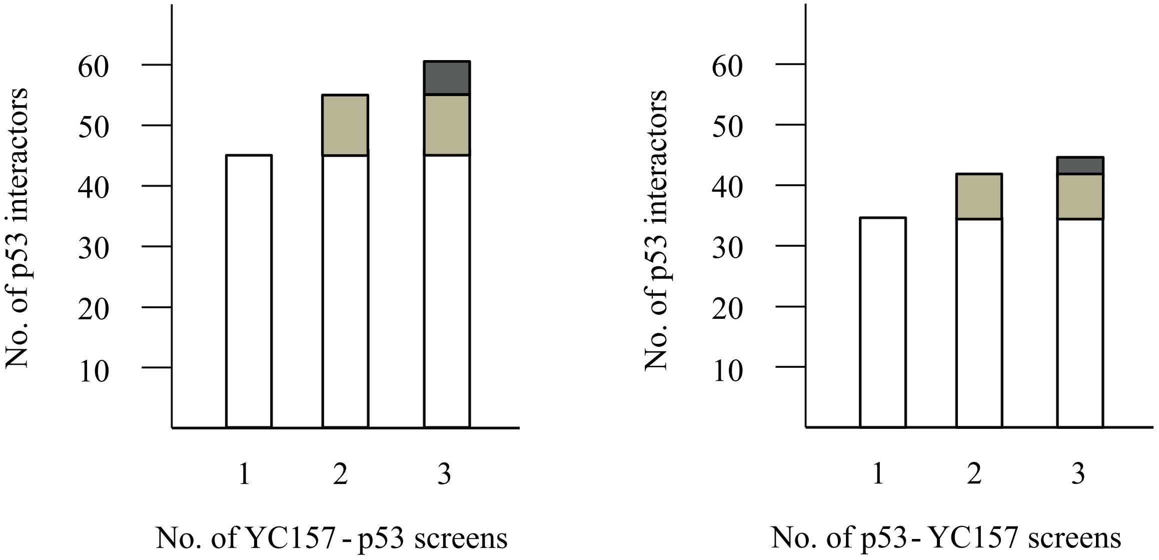


**Supplementary figure 7**

### supplementary figure 8

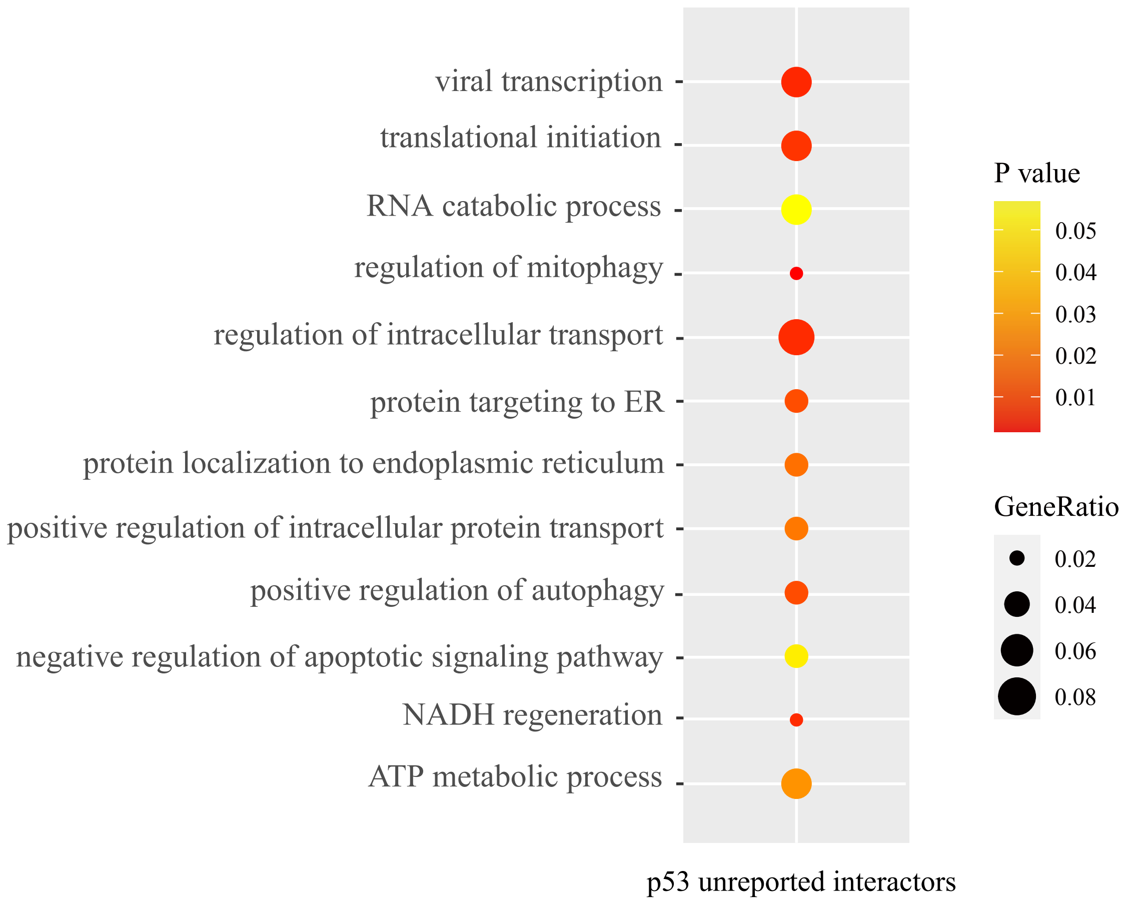


**Supplementary figure 8**

### supplementary figure 9

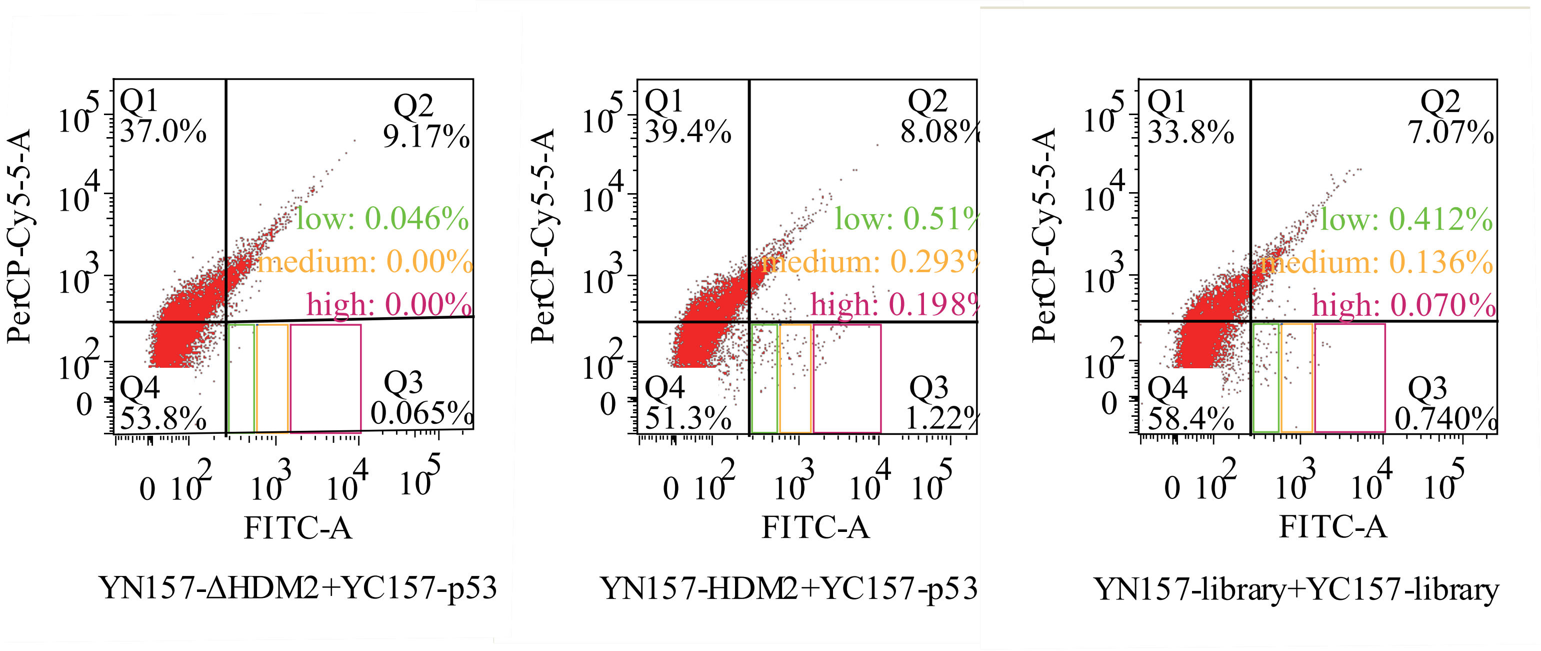


**Supplementary Figure 9**

### supplementary figure 10

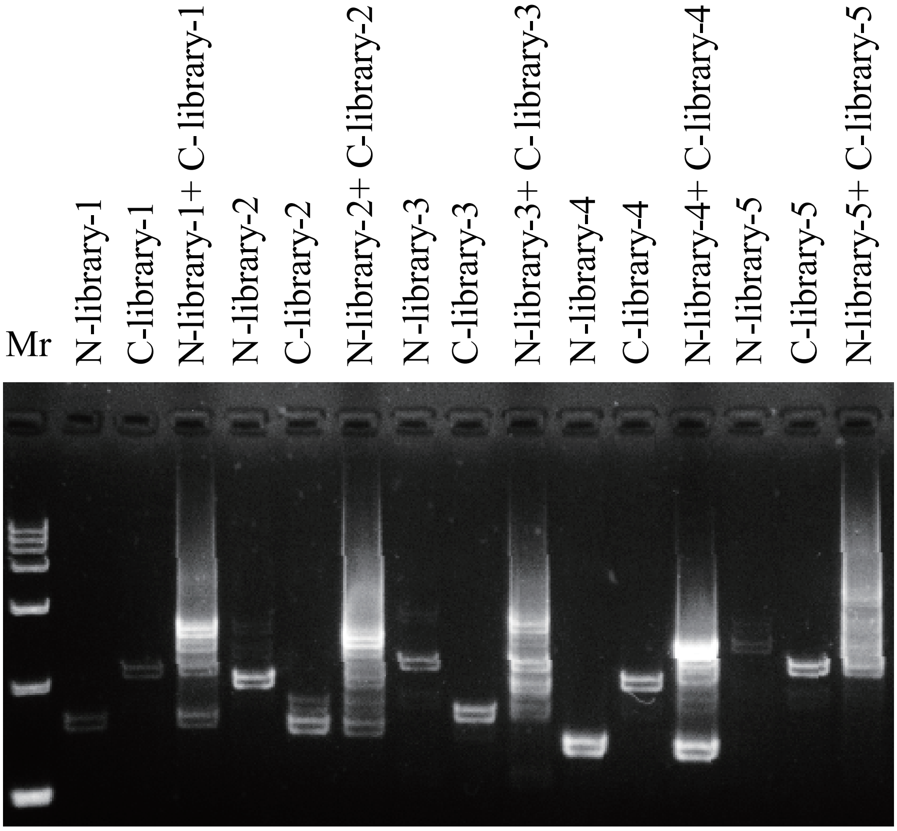


**Supplementary Figure 10**
