## supplementary figure 3 for "A yeast BiFC-seq method for genome-wide interactome mapping"

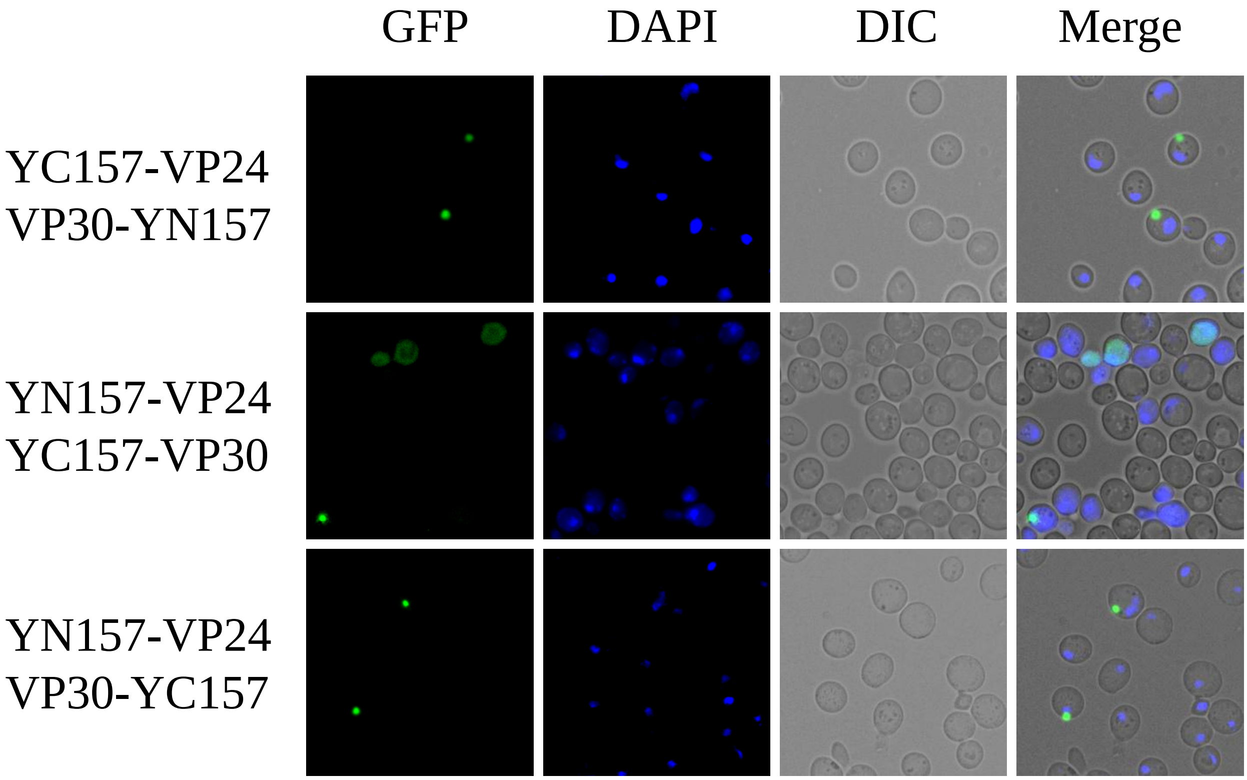


Interaction of VP24 and VP30

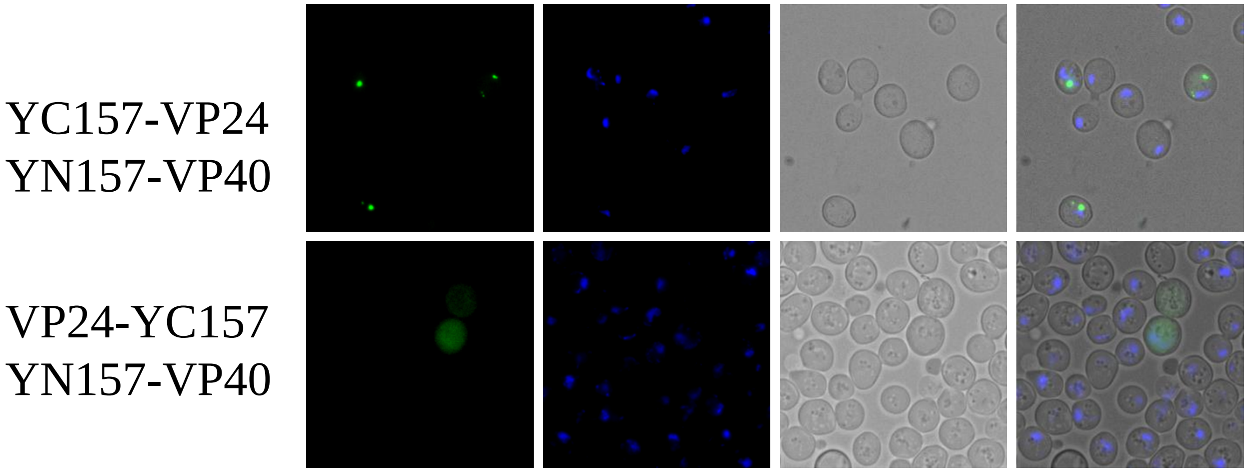


Interaction of VP24 and VP40

YN157-VP24
NP-YC157

YN157-VP24
YC157-NP

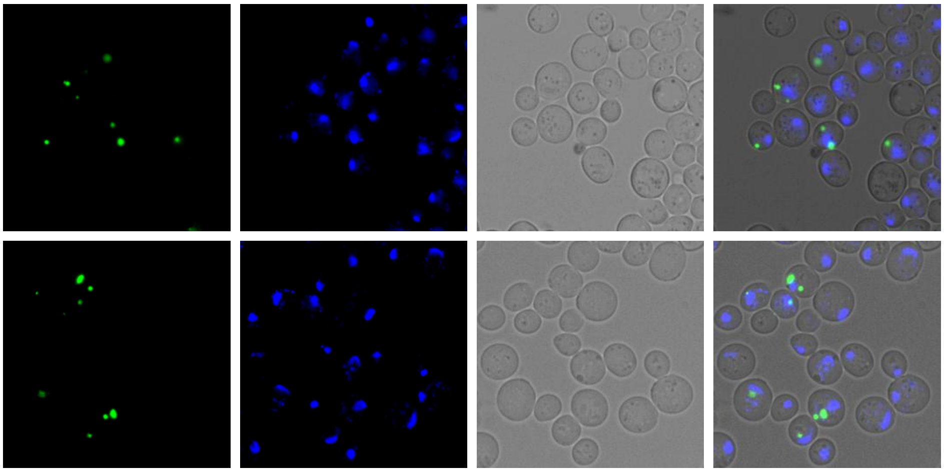

Interaction of VP24 and NP

ssGP-YN157
VP24-YC157

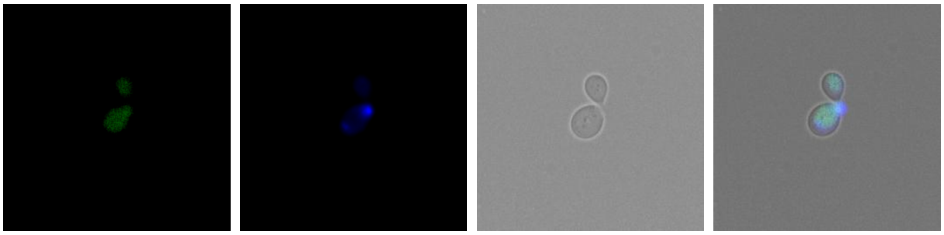

Interaction of VP24 and ssGP

YN157-VP40
YC157-VP40

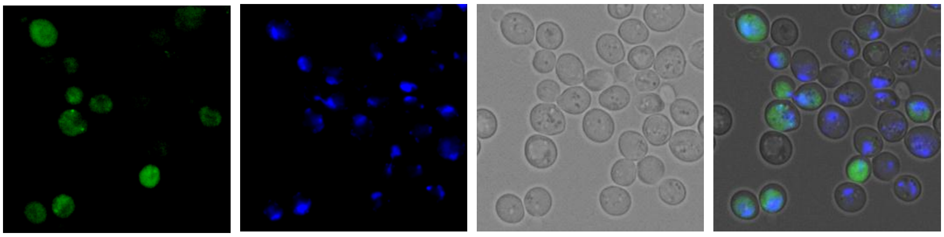

Interaction of VP40 and VP40

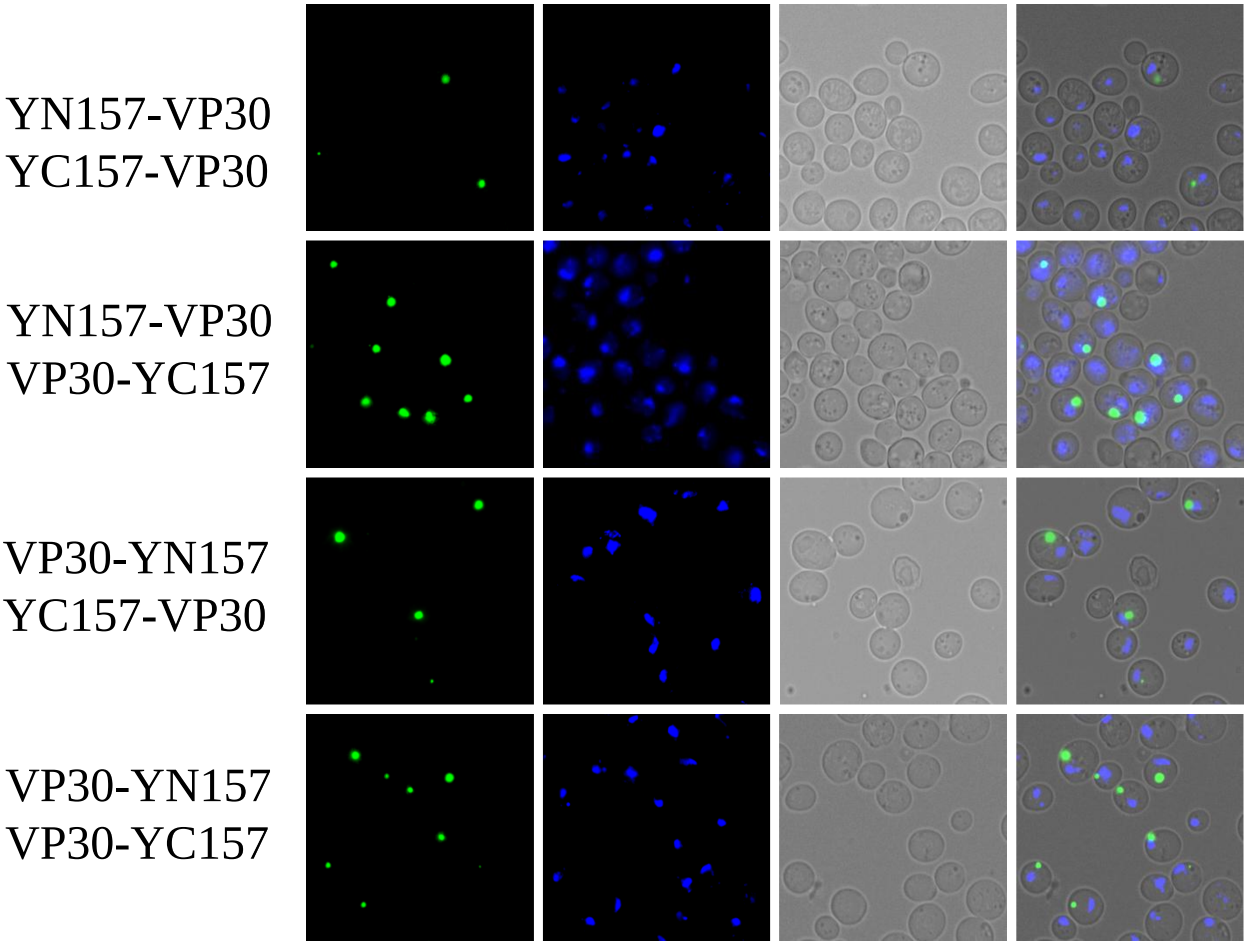


Interaction of VP30 and VP30

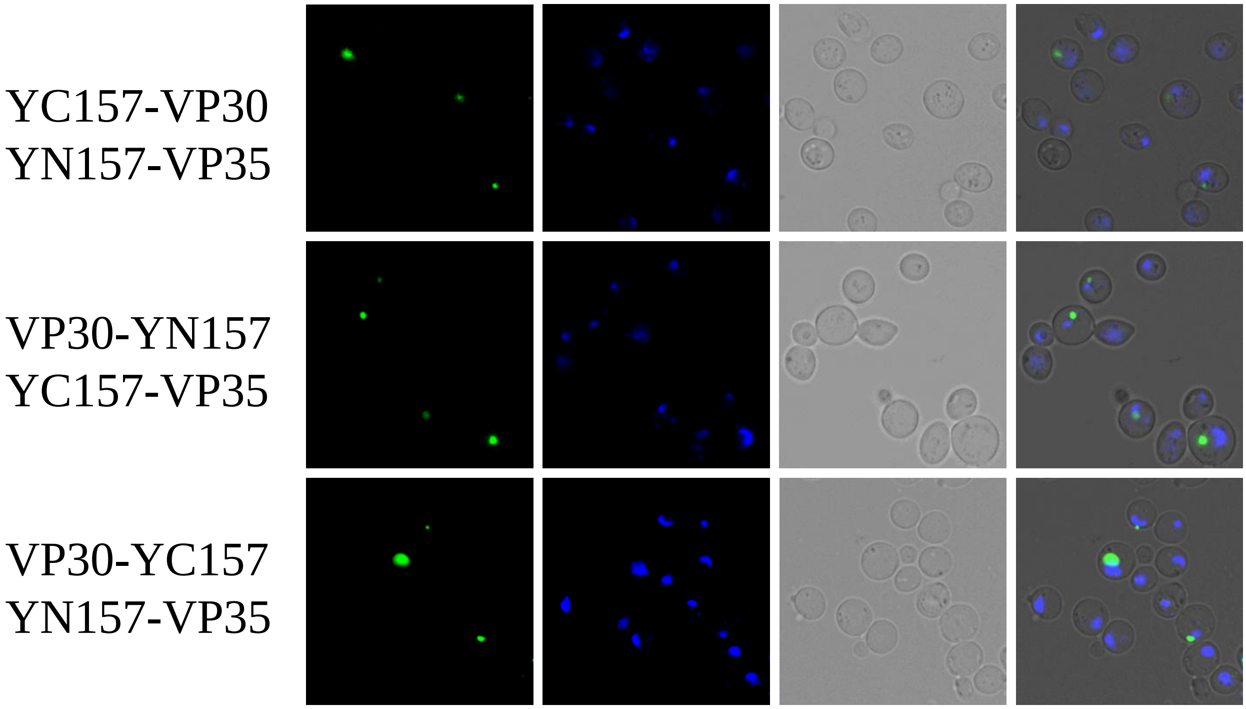


Interaction of VP30 and VP35

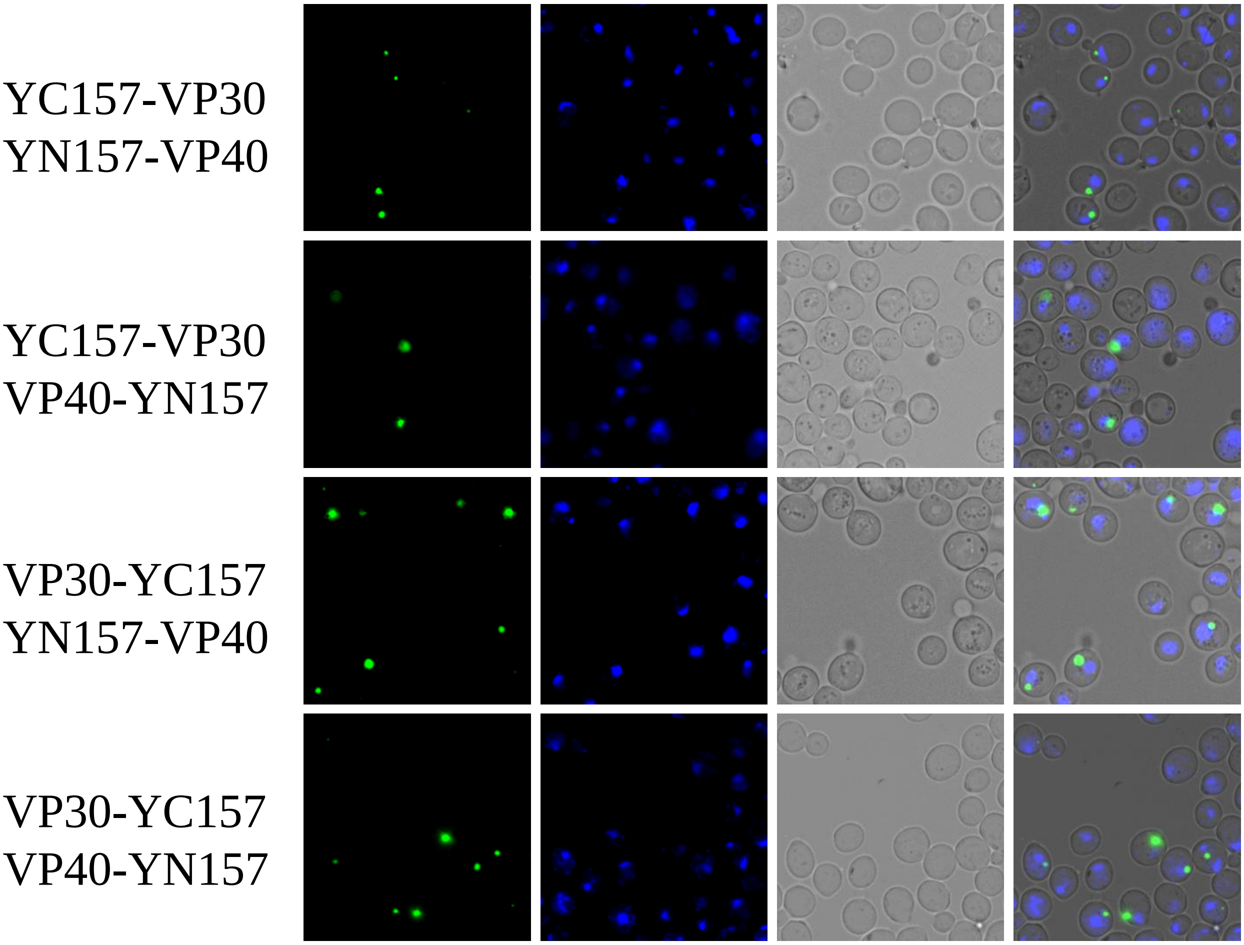


Interaction of VP30 and VP40

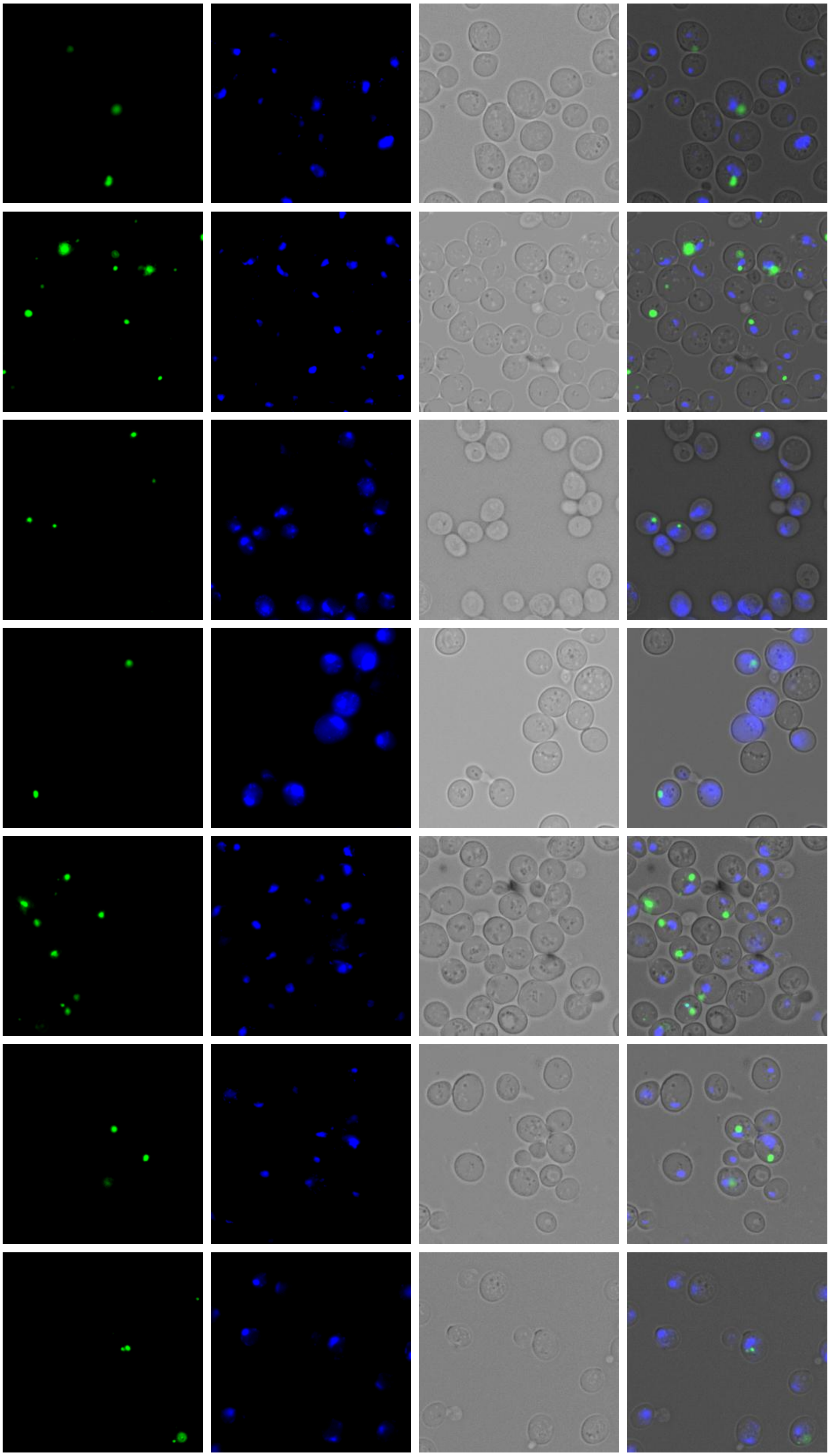


Interaction of VP30 and NP

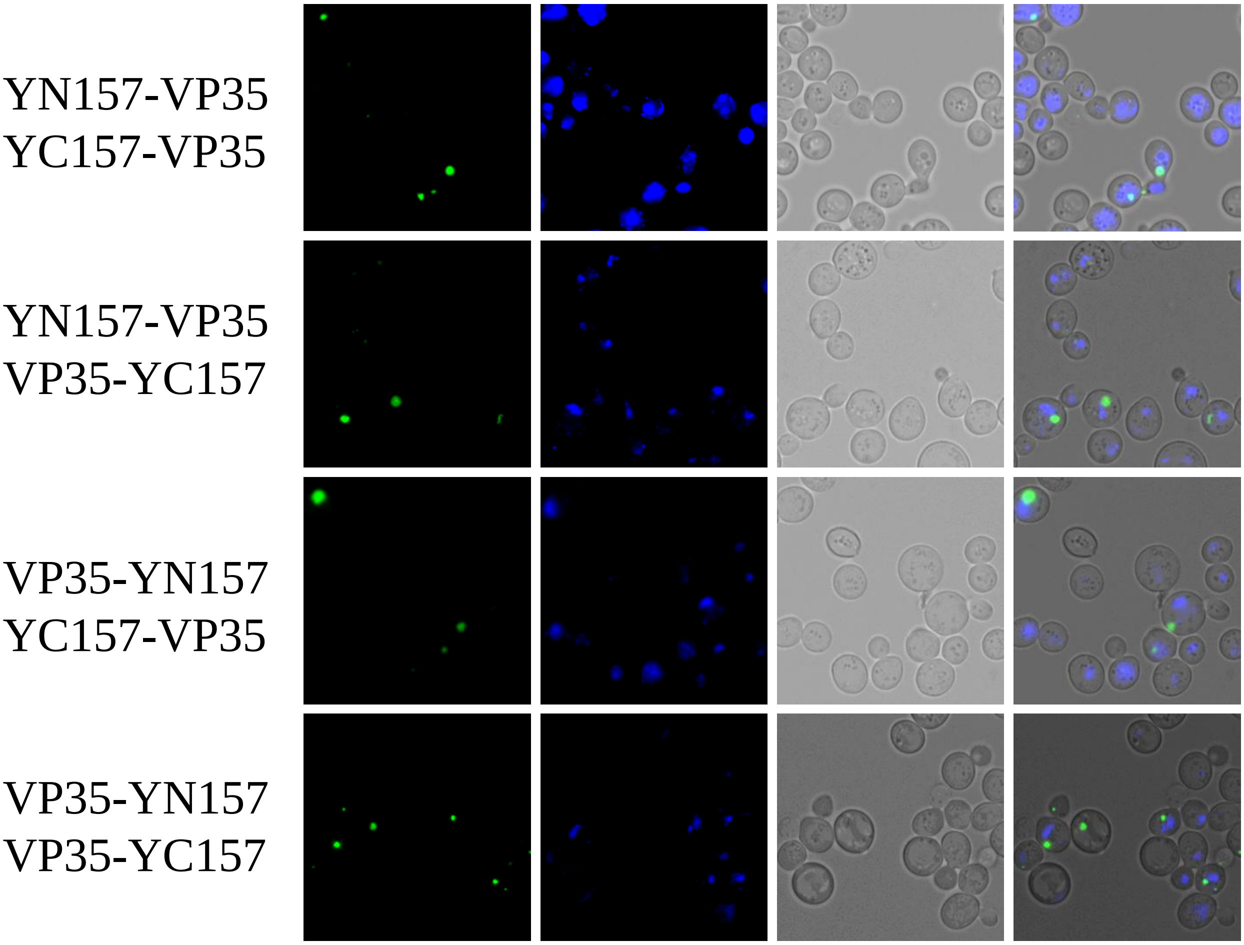


Interaction of VP35 and VP35

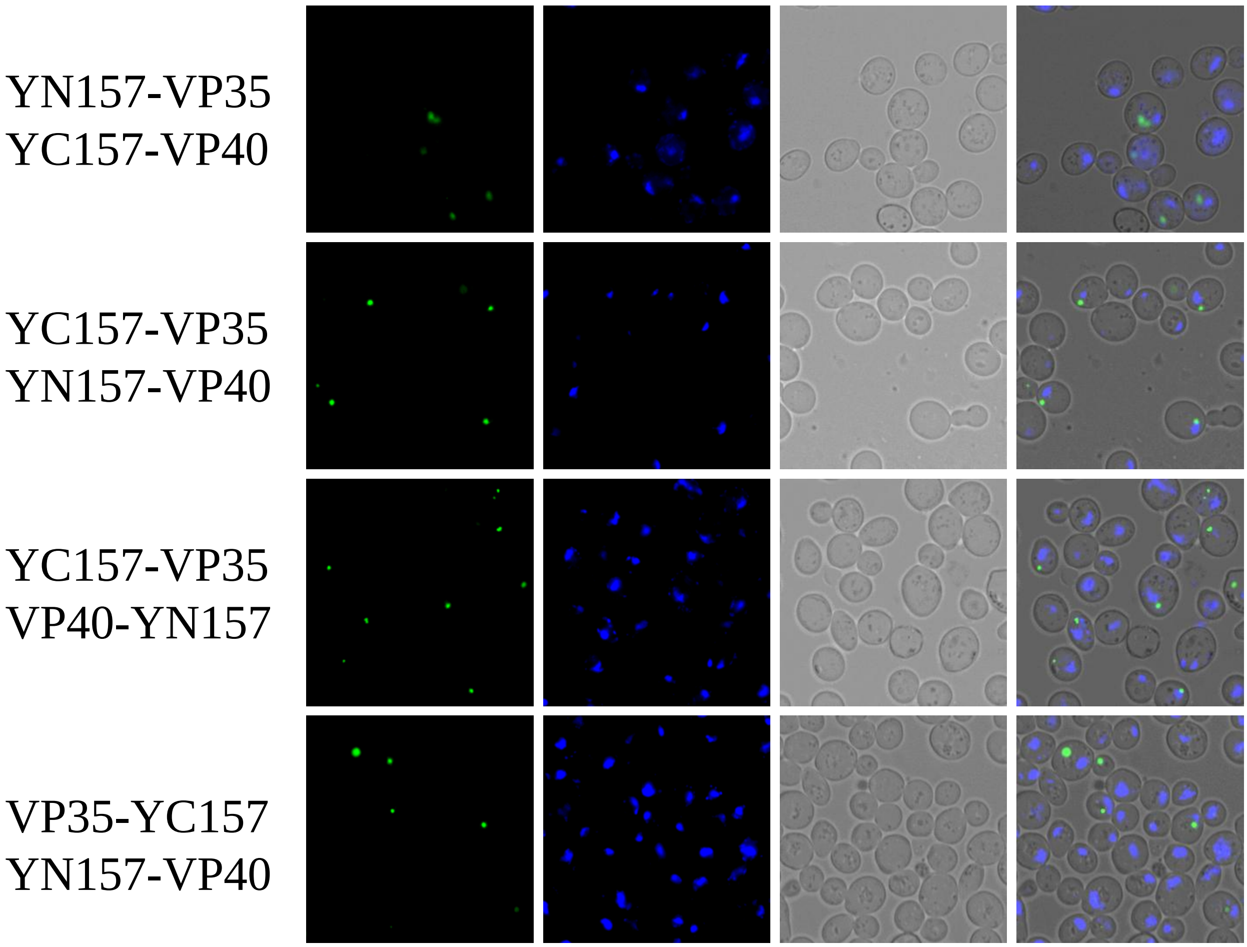


Interaction of VP35 and VP40

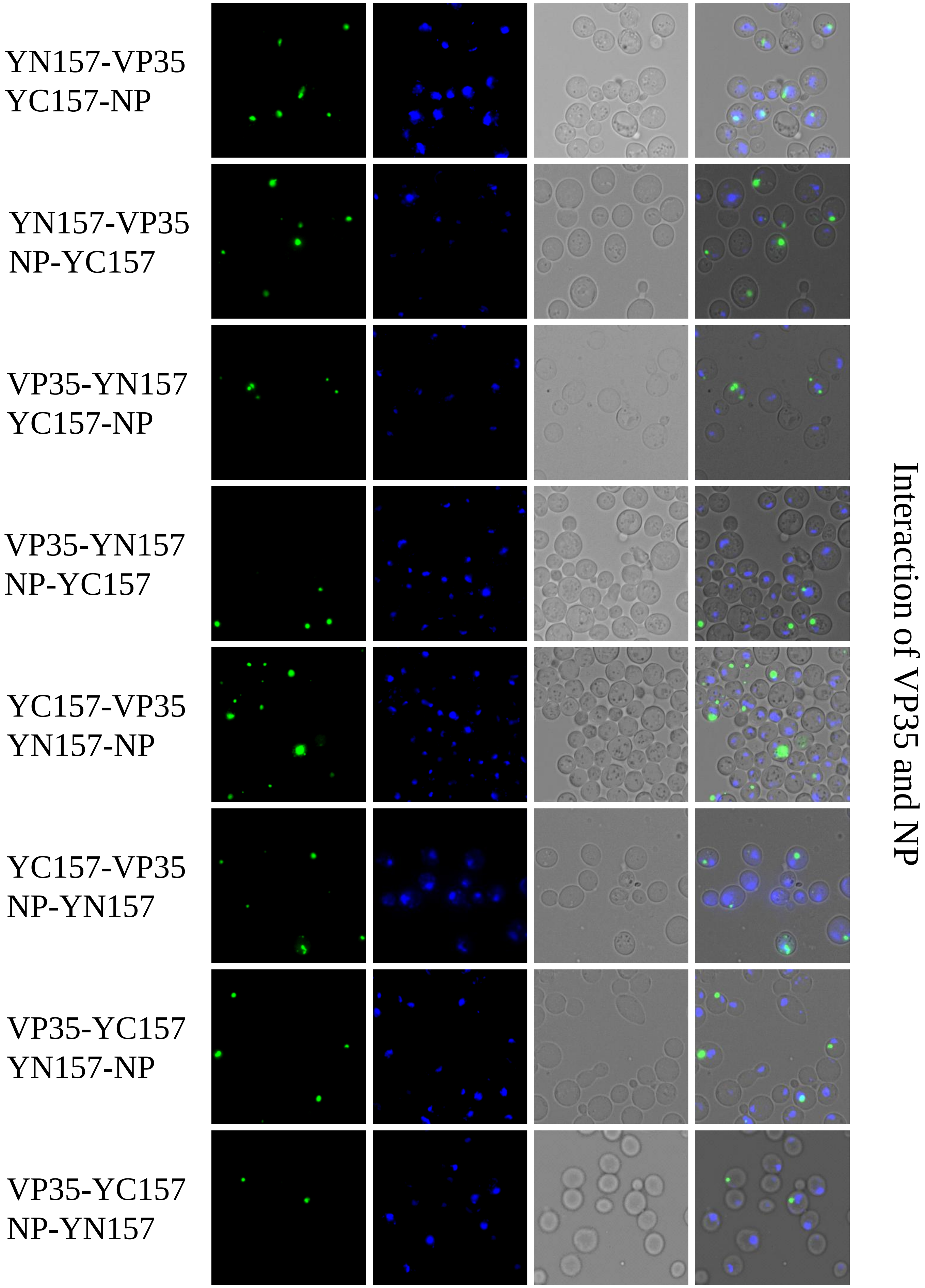
YN157-VP35
YC157-NP
YN157-VP35
NP-YC157
VP35-YN157
YC157-NP
VP35-YN157
NP-YC157
YC157-VP35
YN157-NP
YC157-VP35
NP-YN157
VP35-YC157
YN157-NP
VP35-YC157
NP-YN157
Interaction of VP35 and NP

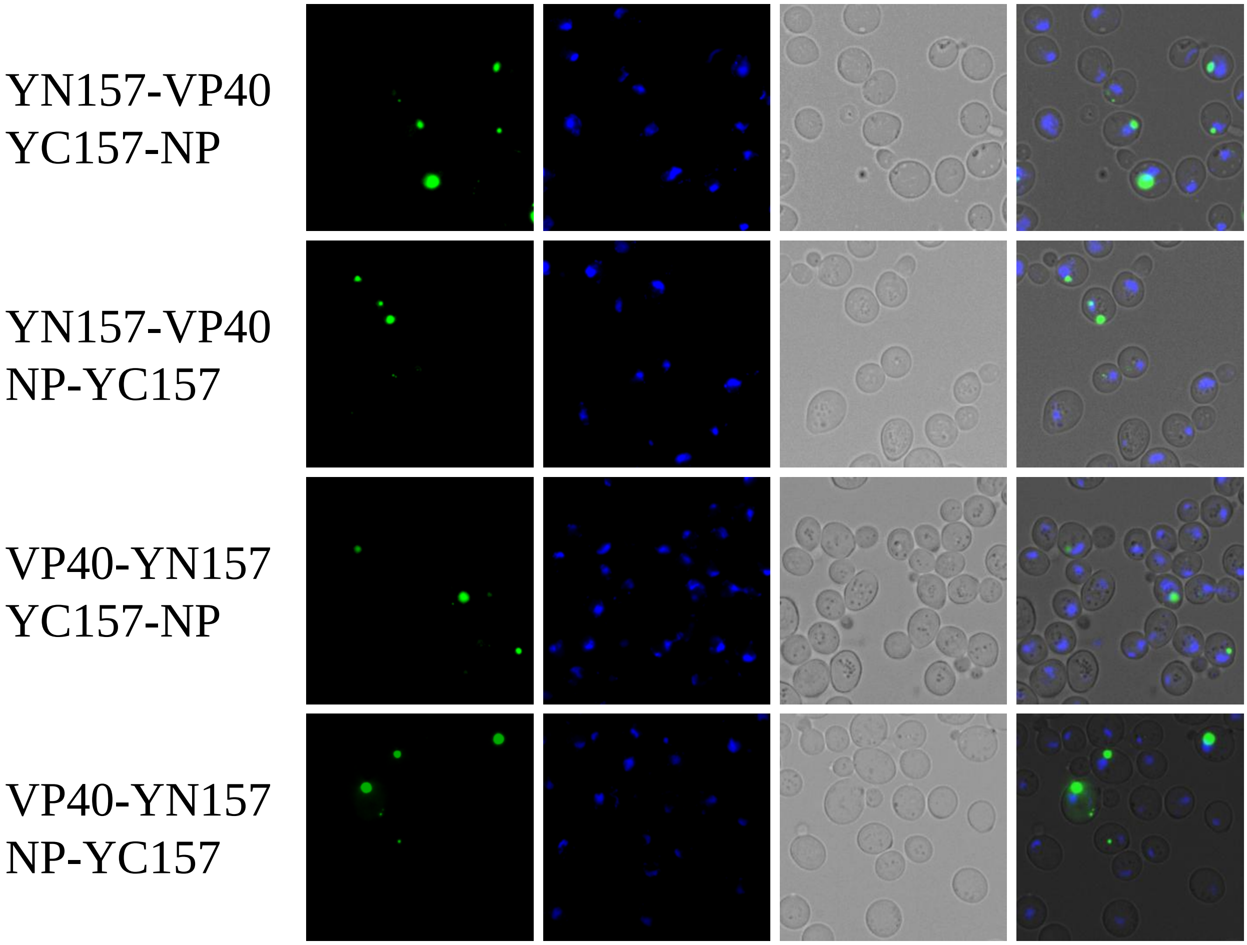


Interaction of VP40 and NP

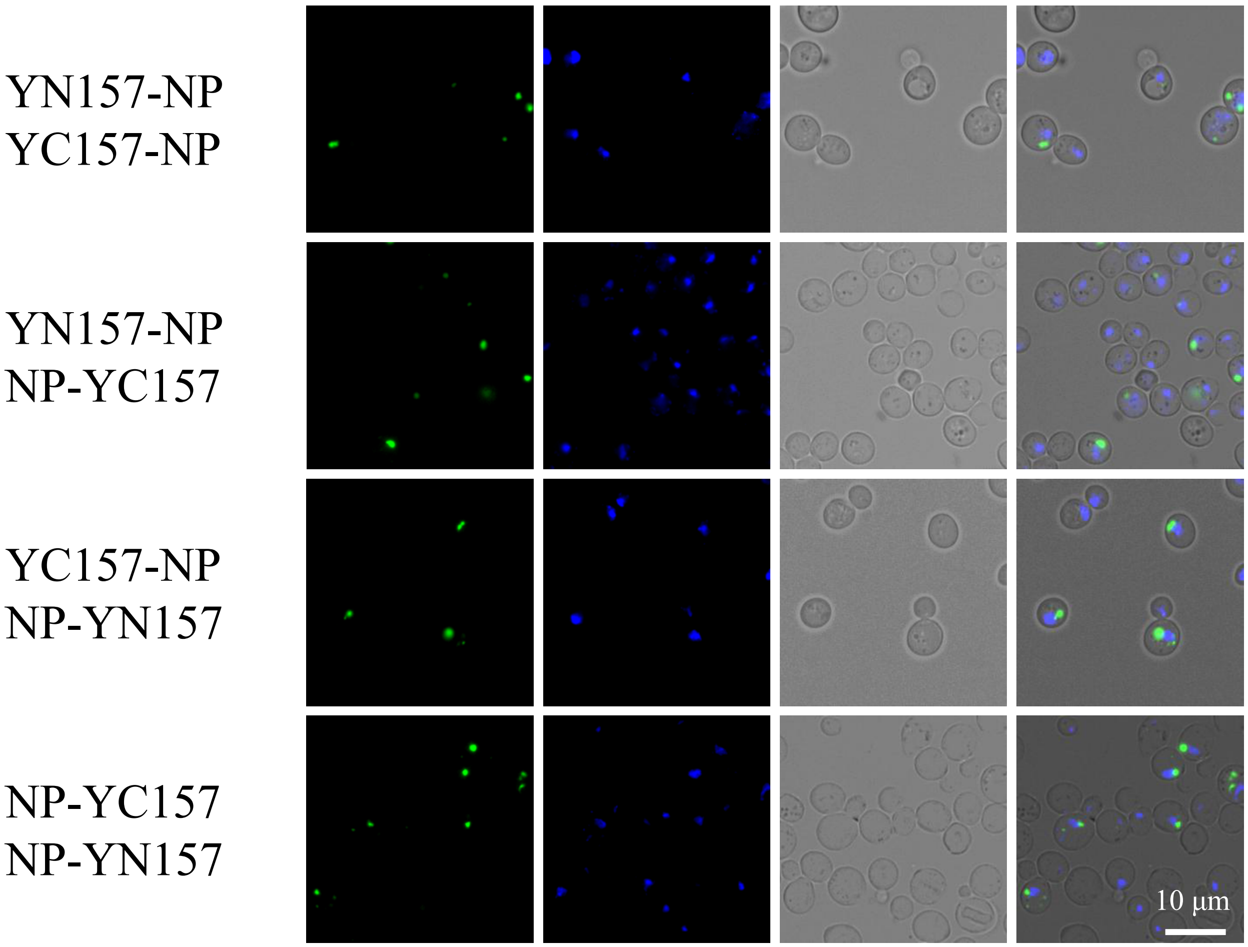


Interaction of NP and NP

Supplementary figure 3
