## supplementary table 1 for "A yeast BiFC-seq method for genome-wide interactome mapping"

| Group | Gene Symbol | Recorded in BioGrid Database |
|---|---|---|
| YC157-p53 | AKAP1 | |
| YC157-p53 | ALDOB | |
| YC157-p53 | ALPL | |
| YC157-p53 | AMZ2 | |
| YC157-p53 | AP2M1 | |
| YC157-p53 | ARL6IP4 | |
| YC157-p53 | C9orf16 | |
| YC157-p53 | CALD1 | Yes |
| YC157-p53 | CCDC74B | |
| YC157-p53 | CCDC96 | |
| YC157-p53 | CMTM2 | |
| YC157-p53 | COL1A2 | |
| YC157-p53 | CORO7 | |
| YC157-p53 | ECHDC2 | |
| YC157-p53 | EGFL7 | |
| YC157-p53 | FAT2 | |
| YC157-p53 | FTL | |
| YC157-p53 | GSN | Yes |
| YC157-p53 | HMGN1 | |
| YC157-p53 | HMOX1 | |
| YC157-p53 | KANK3 | |
| YC157-p53 | MARS | |
| YC157-p53 | MBOAT7 | |
| YC157-p53 | MLF2 | |
| YC157-p53 | NDUFS6 | |
| YC157-p53 | NDUFS8 | |
| YC157-p53 | NSFL1C | |
| YC157-p53 | OAZ1 | |
| YC157-p53 | OTUB1 | Yes |
| YC157-p53 | PDZD8 | |
| YC157-p53 | PGK1 | |
| YC157-p53 | PLAUR | |
| YC157-p53 | POLD2 | |
| YC157-p53 | PREB | |
| YC157-p53 | PRMT1 | Yes |
| YC157-p53 | PRPF3 | |
| YC157-p53 | PSMC5 | Yes |

| Group | Gene Symbol | Recorded in BioGrid Database |
|---|---|---|
| YC157-p53 | RPL18 | Yes |
| YC157-p53 | RPL29 | |
| YC157-p53 | RPS18 | Yes |
| YC157-p53 | RPS20 | |
| YC157-p53 | RPS3 | Yes |
| YC157-p53 | RPS5 | |
| YC157-p53 | RRP1 | Yes |
| YC157-p53 | RUSC2 | |
| YC157-p53 | SAMD4A | |
| YC157-p53 | SIGMAR1 | |
| YC157-p53 | SNRPN | Yes |
| YC157-p53 | TAGLN | |
| YC157-p53 | TMSB4X | Yes |
| YC157-p53 | TOMM7 | |
| YC157-p53 | TSC2 | |
| YC157-p53 | ZDHHC12 | |
| YC157-p53 | ZNF365 | |
| p53-YC157 | ABCD4 | |
| p53-YC157 | C16orf13 | |
| p53-YC157 | CAMLG | |
| p53-YC157 | CCDC159 | |
| p53-YC157 | CCT3 | Yes |
| p53-YC157 | CD63 | |
| p53-YC157 | CDK5 | Yes |
| p53-YC157 | CPSF4L | |
| p53-YC157 | CRIP2 | |
| p53-YC157 | CRYAB | Yes |
| p53-YC157 | DTNBP1 | |
| p53-YC157 | FBXW5 | |
| p53-YC157 | GTF3C6 | |
| p53-YC157 | GZF1 | |
| p53-YC157 | HLA-DMB | |
| p53-YC157 | IFI6 | |
| p53-YC157 | IFITM3 | |
| p53-YC157 | IGLL1 | |
| p53-YC157 | JUP | |
| p53-YC157 | KLHDC9 | |
| p53-YC157 | MPC2 | |
| p53-YC157 | MPLKIP | |
| p53-YC157 | NCAPH2 | |
| p53-YC157 | NDN | Yes |
| p53-YC157 | NECAB2 | |

| Group | Gene Symbol | Recorded in BioGrid Database |
|---|---|---|
| p53-YC157 | NEDD9 | |
| p53-YC157 | NFKBIA | Yes |
| p53-YC157 | PDZK1IP1 | |
| p53-YC157 | RPL28 | Yes |
| p53-YC157 | RPL7A | Yes |
| p53-YC157 | SCAMP3 | |
| p53-YC157 | SLIT3 | |
| p53-YC157 | SUPT7L | Yes |
| p53-YC157 | TCTN3 | |
| p53-YC157 | TXNDC2 | |
| p53-YC157 | YWHAZ | Yes |
| p53 | ARRDC5 | |
| p53 | EIF4G2 | |
| p53 | MSRA | |
| p53 | PPM1G | |
| p53 | RSRP1 | |
| p53 | SPATS2 | |
| p53 | TPI1 | Yes |

**Supplementary Table 1**
