## supplementary table 3 for "A yeast BiFC-seq method for genome-wide interactome mapping"

| Evidence | Interactions Submitted | High Confidence, LR>2.0 | PPI Database | Interolog | Interacting Domain | GO Coannotation | Genome Context | Gene Coexpression | Network Topology |
|---|---|---|---|---|---|---|---|---|---|
| #interaction | 53 | 34 | 0 | 0 | 8 | 34 | 0 | 11 | 10 |

**Supplementary Table 2**
