## supplementary table 3 for "A yeast BiFC-seq method for genome-wide interactome mapping"

| GO ID | Description | Gene Ratio | BgRatio | p value | p adjust | q value | geneID | Count |
|---|---|---|---|---|---|---|---|---|
| GO:0006613 | cotranslational protein targeting to membrane | 8/85 | 110/18679 | 3.69E-08 | 1.42E-05 | 1.20E-05 | 6130/6141/6158/6159/6188/6193/6222/6224 | 8 |
| GO:0045047 | protein targeting to ER | 8/85 | 114/18679 | 4.89E-08 | 1.42E-05 | 1.20E-05 | 6130/6141/6158/6159/6188/6193/6222/6224 | 8 |
| GO:0070972 | protein localization to endoplasmic reticulum | 8/85 | 137/18679 | 2.04E-07 | 2.83E-05 | 2.39E-05 | 6130/6141/6158/6159/6188/6193/6222/6224 | 8 |
| GO:0000956 | nuclea-rtranscribed mRNA catabolic process | 8/85 | 191/18679 | 2.54E-06 | 0.000177 | 0.00015 | 6130/6141/6158/6159/6188/6193/6222/6224 | 8 |
| GO:0006413 | translational initiation | 9/85 | 262/18679 | 2.98E-06 | 0.000201 | 0.00017 | 1982/6130/6141/6158/6159/6188/6193/6222/622 | 49 |
| GO:0006414 | translational elongation | 8/85 | 204/18679 | 4.13E-06 | 0.000247 | 0.000209 | 6130/6141/6158/6159/6188/6193/6222/6224 | 8 |
| GO:0006401 | RNA catabolic process | 8/85 | 234/18679 | 1.13E-05 | 0.000577 | 0.000488 | 6130/6141/6158/6159/6188/6193/6222/6224 | 8 |
| GO:0032507 | maintenance of protein location in cell | 5/85 | 92/18679 | 6.24E-05 | 0.002609 | 0.00221 | 2934/3728/4792/7114/9913 | 5 |
| GO:0044270 | cellular nitrogen compound catabolic process | 9/85 | 440/18679 | 0.000174 | 0.006057 | 0.00513 | 3162/6130/6141/6158/6159/6188/6193/6222/622 | 49 |
| GO:0032387 | negative regulation of intracellular transport | 5/85 | 133/18679 | 0.000353 | 0.011167 | 0.009459 | 1020/1410/3162/4792/8165 | 5 |
| GO:0008347 | glial cell migration | 3/85 | 38/18679 | 0.000684 | 0.019568 | 0.016575 | 1020/4692/79143 | 3 |
| GO:0042989 | sequestering of actin monomer | s 2/85 | 10/18679 | 0.000899 | 0.02505 | 0.021218 | 2934/7114 | 2 |
| GO:0045738 | negative regulation of DNA repair | 2/85 | 11/18679 | 0.001096 | 0.029352 | 0.024862 | 55611/6188 | 2 |
| GO:2001021 | negative regulation of response to DNA damage stimulus | 3/85 | 48/18679 | 0.001357 | 0.03543 | 0.030011 | 55611/6188/7178 | 3 |
| GO:0033157 | regulation of intracellular protein transport | 7/85 | 366/18679 | 0.001395 | 0.035967 | 0.030466 | 1020/3728/4792/4946/54543/6158/8165 | 7 |
| GO:2001233 | regulation of apoptotic signaling pathway | 7/85 | 380/18679 | 0.001727 | 0.04296 | 0.036389 | 2537/2934/3162/5329/6188/7178/7534 | 7 |

**Supplementary Table3**
