## supplementary table 4 for "A yeast BiFC-seq method for genome-wide interactome mapping"

| GO ID | Description | GeneRatio | BgRatio | pvalue | geneID |
|---|---|---|---|---|---|
| GO:0032386 | regulation of intracellular transport | 6/67 | 423/18670 | 0.004374509 | 3162/4946/10113/54543/84062/3728 |
| GO:0006413 | translational initiation | 4/67 | 193/18670 | 0.005385045 | 6159/6224/6193/1982 |
| GO:0045047 | protein targeting to ER | 3/67 | 118/18670 | 0.00915287 | 6159/6224/6193 |
| GO:0090316 | positive regulation of intracellular protein transport | 3/67 | 153/18670 | 0.018345708 | 4946/54543/3728 |
| GO:2001234 | negative regulation of apoptotic signaling pathway | 3/67 | 230/18670 | 0.051577168 | 3162/5329/2537 |
| GO:0006401 | RNA catabolic process | 4/67 | 397/18670 | 0.056721195 | 6159/6224/6193/23034 |
| GO:0006735 | NADH regeneration | 2/67 | 27/18670 | 0.004325972 | 229/5230 |
| GO:0019083 | viral transcription | 4/67 | 177/18670 | 0.003966733 | 6159/6224/6193/10410 |
| GO:0010508 | positive regulation of autophagy | 3/67 | 117/18670 | 0.008943154 | 3162/54543/7249 |
| GO:0046034 | ATP metabolic process | 4/67 | 305/18670 | 0.025132456 | 229/4726/4728/5230 |
| GO:1901524 | regulation of mitophagy | 2/67 | 17/18670 | 0.001715981 | 54543/7249 |
| GO:0070972 | protein localization to endoplasmic reticulum | 3/67 | 147/18670 | 0.016509467 | 6159/6224/6193 |

**Supplementary Table 4**
