## supplementary table 5 for "A yeast BiFC-seq method for genome-wide interactome mapping"

| **Official Symbol_a** | **Gene ID_a** | **Official Symbol_b** | **Gene ID_b** | **Source** |
|---|---|---|---|---|
| CDKN2C | 1031 | TP53 | 7157 | Cell 122, 957–968, 2005 |
| ERH | 2079 | TP53 | 7157 | Cell 122, 957–968, 2005 |
| ZNF24 | 7572 | TP53 | 7157 | Cell 122, 957–968, 2005 |
| ARIH2 | 10425 | TP53 | 7157 | Cell 122, 957–968, 2005 |
| BTBD2 | 55643 | TP53 | 7157 | Cell 122, 957–968, 2005 |
| CCL18 | 6362 | TP53 | 7157 | Cell 122, 957–968, 2005 |
| CDC42 | 998 | TP53 | 7157 | Cell 122, 957–968, 2005 |
| EIF2S2 | 8894 | TP53 | 7157 | Cell 122, 957–968, 2005 |
| LAMA4 | 3910 | TP53 | 7157 | Cell 122, 957–968, 2005 |
| MAD2L1BP | 9587 | TP53 | 7157 | Cell 122, 957–968, 2005 |
| MPHOSPH6 | 10200 | TP53 | 7157 | Cell 122, 957–968, 2005 |
| PAFAH1B3 | 5050 | TP53 | 7157 | Cell 122, 957–968, 2005 |
| PCDHA4 | 56144 | TP53 | 7157 | Cell 122, 957–968, 2005 |
| PP | 5464 | TP53 | 7157 | Cell 122, 957–968, 2005 |
| PSMD11 | 5717 | TP53 | 7157 | Cell 122, 957–968, 2005 |
| SAT | 6303 | TP53 | 7157 | Cell 122, 957–968, 2005 |
| SERPINB9 | 5272 | TP53 | 7157 | Cell 122, 957–968, 2005 |
| SMA3 | 10571 | TP53 | 7157 | Cell 122, 957–968, 2005 |
| SNRPN | 6638 | TP53 | 7157 | Cell 122, 957–968, 2005 |
| SULT1E1 | 6783 | TP53 | 7157 | Cell 122, 957–968, 2005 |
| TK1 | 7083 | TP53 | 7157 | Cell 122, 957–968, 2005 |
| Tmsb4x | 19241 | TP53 | 7157 | Cell 122, 957–968, 2005 |
| ANXA3 | 306 | TP53 | 7157 | Cell 122, 957–968, 2005 |
| ARL3 | 403 | TP53 | 7157 | Cell 122, 957–968, 2005 |
| ATF3 | 467 | TP53 | 7157 | Cell 122, 957–968, 2005 |
| BCR | 613 | TP53 | 7157 | Cell 122, 957–968, 2005 |
| CCT5 | 22948 | TP53 | 7157 | Cell 122, 957–968, 2005 |
| COX17 | 10063 | TP53 | 7157 | Cell 122, 957–968, 2005 |
| DLEU1 | 10301 | TP53 | 7157 | Cell 122, 957–968, 2005 |
| FXYD6 | 53826 | TP53 | 7157 | Cell 122, 957–968, 2005 |
| GSTM4 | 2948 | TP53 | 7157 | Cell 122, 957–968, 2005 |
| HSPB1 | 3315 | TP53 | 7157 | Cell 122, 957–968, 2005 |
| NP | 4860 | TP53 | 7157 | Cell 122, 957–968, 2005 |
| STX5A | 6811 | TP53 | 7157 | Cell 122, 957–968, 2005 |
| TP53BP1 | 7158 | TP53 | 22059 | Cell 122, 957–968, 2005 |
| WDR33 | 55339 | TP53 | 7157 | Cell 122, 957–968, 2005 |

| **Official Symbol_a** | **Gene ID_a** | **Official Symbol_b** | **Gene ID_b** | **Source** |
|---|---|---|---|---|
| HSU79303 | 29903 | TP53 | 7157 | Cell 122, 957–968, 2005 |
| MGC2494 | 65990 | TP53 | 7157 | Cell 122, 957–968, 2005 |
| S100A8 | 6279 | TP53 | 7157 | Cell 122, 957–968, 2005 |
| ZCCHC10 | 54819 | TP53 | 7157 | Cell 122, 957–968, 2005 |
| KIAA0087 | 9808 | TP53 | 7157 | Cell 122, 957–968, 2005 |
| RAB4A | 5867 | TP53 | 7157 | Cell 122, 957–968, 2005 |
| THAP8 | 199745 | TP53 | 7157 | Cell 122, 957–968, 2005 |
| DVL2 | 1856 | TP53 | 7157 | Nature 437, 1173–1178, 2005 |
| MGC33889 | 286514 | TP53 | 7157 | Nature 437, 1173–1178, 2005 |
| GPX2 | 2877 | TP53 | 7157 | Mol Syst Biol 7, 536, 2011 |
| MDM2 | 4193 | TP53 | 7157 | Mol Syst Biol 7, 536, 2011 |
| TP53 | 7157 | TP53 | 7157 | Mol Syst Biol 7, 536, 2011 |
| RCHY1 | 25898 | TP53 | 7157 | Mol Syst Biol 7, 536, 2011 |
| SETD7 | 80854 | TP53 | 7157 | Mol Syst Biol 7, 536, 2011 |
| FCAMR | 83953 | TP53 | 7157 | Mol Syst Biol 7, 536, 2011 |

**Supplementary Table 5**
