## supplementary table 6 for "A yeast BiFC-seq method for genome-wide interactome mapping"

| Gene Symbol_a | Gene Symbol_b | Fluorescence intensity | Recorded in BioGrid Database |
|---|---|---|---|
| RPL30 | RPL13 | high | Yes |
| RPL9 | RPL27A | high | Yes |
| VIMP | UFD1L | high | Yes |
| HLA-A | GAP43 | high | |
| ANKMY2 | TPT1 | high | |
| ANKMY2 | APOE | high | |
| ANKMY2 | RPL29 | high | |
| BRD3 | TMBIM6 | high | |
| BRD3 | ALDOA | high | |
| C17orf70 | PABPC1 | high | |
| C4orf3 | CCDC89 | high | |
| C4orf3 | MYLK | high | |
| C4orf3 | HADHA | high | |
| C4orf3 | MGST1 | high | |
| C4orf3 | SHARPIN | high | |
| C4orf3 | MT3 | high | |
| C4orf3 | ARL3 | high | |
| C4orf3 | GAP43 | high | |
| CD81 | PDHA2 | high | |
| CD81 | BFSP1 | high | |
| CERS2 | ALDOA | high | |
| CMTM2 | TBCB | high | |
| CMTM2 | UBE2S | high | |
| CMTM2 | TPM2 | high | |
| CREG1 | TMSB4X | high | |
| CST3 | RPL23A | high | |
| DAXX | ANXA2 | high | |
| EIF1 | APRT | high | |
| EIF1 | EEF1G | high | |
| FBXW5 | BFSP1 | high | |
| FRG1 | PABPC1 | high | |

| Gene Symbol_a | Gene Symbol_b | Fluorescence intensity | Recorded in BioGrid Database |
|---|---|---|---|
| FXYD2 | CCDC89 | high | |
| GPX4 | RBM34 | high | |
| GTPBP6 | ARHGAP42 | high | |
| GTPBP6 | EEF1G | high | |
| GTPBP6 | RPL31 | high | |
| GTPBP6 | UFD1L | high | |
| H3F3B | SPATA7 | high | |
| HMGCL | UFD1L | high | |
| HMGCL | PABPC1 | high | |
| KAT8 | CCL2 | high | |
| MBP | EEF1G | high | |
| MT1F | PSMC3 | high | |
| MT1F | TMEM223 | high | |
| NME2 | TMBIM6 | high | |
| NPIPB5 | AKR7A3 | high | |
| NPIPB5 | PRM2 | high | |
| OTUB1 | TSSC4 | high | |
| OTUB1 | ATXN2L | high | |
| OTUB1 | REPIN1 | high | |
| OTUB1 | EEF1G | high | |
| OTUB1 | CCDC89 | high | |
| PDZK1IP1 | GTF3C1 | high | |
| PDZK1IP1 | HADHA | high | |
| PFDN5 | APOE | high | |
| PFDN5 | VAMP2 | high | |
| PFDN5 | RPL13 | high | |
| PFDN5 | ALDOB | high | |
| PFDN5 | RPL31 | high | |
| PNPLA6 | PCBP1 | high | |
| PRM2 | CCDC89 | high | |
| PRM3 | HLA-C | high | |
| PRPF6 | TMSB10 | high | |
| PSMD8 | RPL31 | high | |
| RBP1 | CD99 | high | |
| RBP1 | PEBP1 | high | |
| RBP1 | PABPC1 | high | |
| RPL23A | VDAC1 | high | |
| RPL23A | CSDE1 | high | |
| RPL23A | LRRC48 | high | |
| RPL29 | ARL3 | high | |
| RPL29 | DHFR | high | |
| RPL8 | REPIN1 | high | |

| Gene Symbol_a | Gene Symbol_b | Fluorescence intensity | Recorded in BioGrid Database |
|---|---|---|---|
| RPS11 | CST3 | high | |
| RPS19 | CCDC89 | high | |
| RPS19 | PUF60 | high | |
| RPS20 | CIRBP | high | |
| S100A9 | APOE | high | |
| SDSL | UFD1L | high | |
| SFTPC | AKR7A3 | high | |
| SLCO3A1 | MIOX | high | |
| STMN1 | CCDC89 | high | |
| TIGD1 | MGST1 | high | |
| TIGD1 | MYLK | high | |
| TIGD1 | APOE | high | |
| TPM2 | KRT18 | high | |
| TREX1 | TMEM109 | high | |
| TRIM28 | ARL3 | high | |
| TSPAN16 | MT1G | high | |
| TXNDC2 | PSMD10 | high | |
| UBIAD1 | ARHGAP42 | high | |
| UBIAD1 | MT3 | high | |
| VIMP | CCDC89 | high | |
| VIMP | OBSL1 | high | |
| VIMP | RPL23A | high | |
| VIMP | NHP2L1 | high | |
| VIMP | CCL2 | high | |
| VIMP | ALDOA | high | |
| VWF | SPATA7 | high | |
| ZNF365 | BFSP1 | high | |
| HMGN1 | AKR1C3 | high | |
| PDZK1IP1 | RPL11 | high | |
| YBX1 | C1QTNF2 | high | |
| YBX1 | RPL31 | high | |
| ANAPC13 | EEF1G | high | |
| ANXA6 | PGAM2 | high | |
| RPL30 | EEF1G | high | |
| ANXA6 | RPL31 | high | |
| FXYD2 | ALDOA | high | |
| RPS13 | PGS1 | high | |
| RPS11 | PTGES3L-AARSD1 | high | |
| SFTPC | CALM3 | high | |
| OTUB1 | PRDX1 | medium | Yes |
| RPL30 | RPS5 | medium | Yes |
| ANKMY2 | SH3RF2 | medium | |

| Gene Symbol_a | Gene Symbol_b | Fluorescence intensity | Recorded in BioGrid Database |
|---|---|---|---|
| ANKMY2 | TPT1 | medium | |
| APOE | EEF1G | medium | |
| APOE | EIF1 | medium | |
| ARL2 | MT1G | medium | |
| ATP1A1 | TBRG4 | medium | |
| B3GALT6 | GSTO1 | medium | |
| BRD3 | TMBIM6 | medium | |
| BRD3 | CCDC89 | medium | |
| CD74 | ATP6AP1 | medium | |
| CERS2 | RPL35 | medium | |
| CERS2 | RPL30 | medium | |
| COX8A | RPL11 | medium | |
| COX8A | PSMC3 | medium | |
| DHRS4 | SLC25A3 | medium | |
| EEF1A1 | AARSD1 | medium | |
| FCGRT | TMSB4X | medium | |
| FXYD2 | RPL31 | medium | |
| HMGCL | UFD1L | medium | |
| HMGN1 | SORD | medium | |
| KRT19 | PRDX5 | medium | |
| MBP | EEF1G | medium | |
| MBP | RPL31 | medium | |
| MORN3 | TMSB4X | medium | |
| MT1F | TMEM223 | medium | |
| MVD | TARBP2 | medium | |
| OTUB1 | TSSC4 | medium | |
| PCK1 | CSTA | medium | |
| PNPLA6 | PCBP1 | medium | |
| PRM2 | CCDC89 | medium | |
| PRR13 | TMBIM6 | medium | |
| PSAP | LAD1 | medium | |
| PSMB6 | SNRPB2 | medium | |
| RHPN1 | CCS | medium | |
| RPL19 | PSMC3 | medium | |
| RPL23A | ATP5G2 | medium | |
| RPL23A | COX6B1 | medium | |
| RPL30 | CCDC89 | medium | |
| RPL30 | CHKB | medium | |
| RPL7A | EEF1G | medium | |
| RPL9 | EEF1G | medium | |
| RPS19 | PUF60 | medium | |
| SMG6 | CCZ1B | medium | |

| Gene Symbol_a | Gene Symbol_b | Fluorescence intensity | Recorded in BioGrid Database |
|---|---|---|---|
| TREX1 | TMEM109 | medium | |
| VIMP | RPS3A | medium | |
| VWA9 | DCXR | medium | |
| RPL23A | RPL31 | low | Yes |
| RPL23A | GOSR1 | low | Yes |
| RPL23A | RPS3A | low | Yes |
| RPL9 | RPL31 | low | Yes |
| RPS11 | RPL31 | low | Yes |
| RPS13 | RPL31 | low | Yes |
| RPL13 | RPL31 | low | Yes |
| ALDH1A1 | RPL31 | low | |
| ALKBH7 | RPL31 | low | |
| APOE | RPL31 | low | |
| APOE | HOPX | low | |
| APOE | ARL3 | low | |
| ARL2 | MT1G | low | |
| ARL2 | RPL31 | low | |
| BRD3 | TMBIM6 | low | |
| EEFSEC | SLC27A6 | low | |
| EEFSEC | PTGES3L-AARSD1 | low | |
| EIF4A1 | TPT1 | low | |
| FKBP6 | HOPX | low | |
| FXYD2 | RPL31 | low | |
| FXYD2 | WDR59 | low | |
| FXYD2 | PTGES3L-AARSD1 | low | |
| FXYD2 | RPL14 | low | |
| FXYD2 | MST1L | low | |
| FXYD2 | ARL3 | low | |
| FXYD2 | RPL26 | low | |
| FXYD2 | HOPX | low | |
| FXYD2 | GSTP1 | low | |
| FXYD2 | PIN1 | low | |
| FXYD2 | EEF1G | low | |
| FXYD2 | MT1G | low | |
| FXYD2 | PABPC1 | low | |
| FXYD2 | RPL35 | low | |
| FXYD2 | AARSD1 | low | |
| FXYD2 | CCDC89 | low | |
| FXYD2 | RPL9 | low | |
| FXYD2 | TMBIM6 | low | |
| FXYD2 | MIOX | low | |
| FXYD2 | SLC27A6 | low | |

| Gene Symbol_a | Gene Symbol_b | Fluorescence intensity | Recorded in BioGrid Database |
|---|---|---|---|
| FXYD2 | UBE2V2 | low | |
| FXYD2 | TMSB4X | low | |
| FXYD2 | EIF1 | low | |
| FXYD2 | CSTA | low | |
| HMGN1 | MT3 | low | |
| KNDC1 | ARL3 | low | |
| MAP2 | RPL31 | low | |
| MCM7 | GOSR1 | low | |
| MPC2 | RPL31 | low | |
| MPC2 | GOSR1 | low | |
| MT1F | MST1L | low | |
| NDRG2 | HNRNPC | low | |
| NDRG2 | RPL31 | low | |
| NDRG2 | S100A6 | low | |
| PCK1 | CSTA | low | |
| PDXK | TMSB4X | low | |
| PDZK1IP1 | RPL11 | low | |
| PDZK1IP1 | ARL3 | low | |
| PDZK1IP1 | RPL31 | low | |
| PFDN5 | APOE | low | |
| PFDN5 | C11orf58 | low | |
| PPM1G | HOPX | low | |
| PSMD3 | TMSB4X | low | |
| PTGES3L-AARSD1 | RPL31 | low | |
| RABAC1 | GSTP1 | low | |
| RABAC1 | AASDHPPT | low | |
| RPL10 | MT3 | low | |
| RPL23A | AASDHPPT | low | |
| RPL23A | ARL3 | low | |
| RPL23A | HOPX | low | |
| RPL23A | EXOSC5 | low | |
| RPL23A | RPL26 | low | |
| RPL23A | COX6B1 | low | |
| RPL23A | CLU | low | |
| RPS11 | PTGES3L-AARSD1 | low | |
| RPS11 | PIN1 | low | |
| RPS11 | SLC27A6 | low | |
| RPS11 | IAH1 | low | |
| RPS11 | HOPX | low | |
| RPS11 | MT3 | low | |
| RPS11 | LOC81691 | low | |
| RPS11 | AASDHPPT | low | |

| Gene Symbol_a | Gene Symbol_b | Fluorescence intensity | Recorded in BioGrid Database |
|---|---|---|---|
| RPS11 | ANKRD39 | low | |
| RPS13 | EIF1 | low | |
| RPS4Y1 | CSTA | low | |
| SAFB2 | RPL31 | low | |
| SMARCA2 | PTGES3L-AARSD1 | low | |
| SNX12 | RPL31 | low | |
| TSACC | TPT1 | low | |

**Supplementary Table 6**
