## supplementary table 7 for "A yeast BiFC-seq method for genome-wide interactome mapping"

| Evidence | Interactions Submitted | High Confidence, LR>2.0 | PPI Database | Interolog | Interacting Domain | GO Coannotation | Genome Context | Gene Coexpression | Network Topology |
|---|---|---|---|---|---|---|---|---|---|
| #interaction | 137 | 26 | 0 | 0 | 14 | 37 | 0 | 19 | 0 |

**Supplementary Table 7**
