## supplementary table 8 for "A yeast BiFC-seq method for genome-wide interactome mapping"

| Official Symbol_a | Gene ID_a | Official Symbol_b | Gene ID_b | Source | Recorded in BioGrid database |
|---|---|---|---|---|---|
| PGBD2 | 267002 | TP53 | 7157 | Cell 145, 787-799, 2011 | |
| FAM98A | 25940 | TP53 | 7157 | Cell 145, 787-799, 2011 | |
| UBE3A | 7337 | TP53 | 7157 | Cell 145, 787-799, 2011 | Yes |
| CUL7 | 9820 | TP53 | 7157 | Cell 145, 787-799, 2011 | Yes |
| CUL9 | 23113 | TP53 | 7157 | Cell 145, 787-799, 2011 | Yes |
| MAP1S | 55201 | TP53 | 7157 | Cell 145, 787-799, 2011 | |
| PPP4R1 | 9989 | TP53 | 7157 | Cell 145, 787-799, 2011 | |
| A2M | 2 | TP53 | 7157 | Cell 145, 787-799, 2011 | |
| C20orf114 | 92747 | TP53 | 7157 | Cell 145, 787-799, 2011 | |
| GSN | 2934 | TP53 | 7157 | Cell 145, 787-799, 2011 | Yes |
| HNRPDL | 9987 | TP53 | 7157 | Cell 145, 787-799, 2011 | |
| HIST1H4D | 8360 | TP53 | 7157 | Cell 145, 787-799, 2011 | |
| HIST3H2A | 92815 | TP53 | 7157 | Cell 145, 787-799, 2011 | |
| CDC37 | 11140 | TP53 | 7157 | Cell 145, 787-799, 2011 | |
| PPP4C | 5531 | TP53 | 7157 | Cell 145, 787-799, 2011 | Yes |
| ALDH16A1 | 126133 | TP53 | 7157 | Cell 145, 787-799, 2011 | |
| DERA | 51071 | TP53 | 7157 | Cell 145, 787-799, 2011 | |
| PTRF | 284119 | TP53 | 7157 | Cell 145, 787-799, 2011 | |
| VPS35 | 55737 | TP53 | 7157 | Cell 145, 787-799, 2011 | |
| YLPM1 | 56252 | TP53 | 7157 | Cell 145, 787-799, 2011 | |
| FAM161A | 84140 | TP53 | 7157 | Cell 145, 787-799, 2011 | |
| ATE1 | 11101 | TP53 | 7157 | Cell 145, 787-799, 2011 | |
| ARHGEF18 | 23370 | TP53 | 7157 | Cell 145, 787-799, 2011 | |
| SERBP1 | 26135 | TP53 | 7157 | Cell 145, 787-799, 2011 | Yes |
| USP9X | 8239 | TP53 | 7157 | Cell 145, 787-799, 2011 | |
| SART3 | 9733 | TP53 | 7157 | Cell 145, 787-799, 2011 | |
| NCOA5 | 57727 | TP53 | 7157 | Cell 145, 787-799, 2011 | |
| PSPC1 | 55269 | TP53 | 7157 | Cell 145, 787-799, 2011 | |
| TAF15 | 8148 | TP53 | 7157 | Cell 145, 787-799, 2011 | |
| TUBA3C | 7278 | TP53 | 7157 | Cell 145, 787-799, 2011 | |
| TUBA1A | 7846 | TP53 | 7157 | Cell 145, 787-799, 2011 | Yes |
| SNRNP40 | 9410 | TP53 | 7157 | Cell 145, 787-799, 2011 | |
| USP39 | 10713 | TP53 | 7157 | Cell 145, 787-799, 2011 | Yes |
| PPP1CA | 5499 | TP53 | 7157 | Cell 145, 787-799, 2011 | Yes |
| PPP1CC | 5501 | TP53 | 7157 | Cell 145, 787-799, 2011 | Yes |
| PPP1CB | 5500 | TP53 | 7157 | Cell 145, 787-799, 2011 | |
| SET | 6418 | TP53 | 7157 | Cell 145, 787-799, 2011 | Yes |

| Official Symbol_a | Gene ID_a | Official Symbol_b | Gene ID_b | Source | Recorded in BioGrid database |
|---|---|---|---|---|---|
| LOC653889 | 653889 | TP53 | 7157 | Cell 145, 787-799, 2011 | |
| CD2BP2 | 10421 | TP53 | 7157 | Cell 145, 787-799, 2011 | |
| EFTUD2 | 9343 | TP53 | 7157 | Cell 145, 787-799, 2011 | |
| PRPF8 | 10594 | TP53 | 7157 | Cell 145, 787-799, 2011 | |
| PRPF6 | 24148 | TP53 | 7157 | Cell 145, 787-799, 2011 | Yes |
| SNRNP200 | 23020 | TP53 | 7157 | Cell 145, 787-799, 2011 | |
| DDX23 | 9416 | TP53 | 7157 | Cell 145, 787-799, 2011 | |
| PRPF31 | 26121 | TP53 | 7157 | Cell 145, 787-799, 2011 | |
| SART1 | 9092 | TP53 | 7157 | Cell 145, 787-799, 2011 | Yes |
| PRPF4 | 9128 | TP53 | 7157 | Cell 145, 787-799, 2011 | |
| PRPF3 | 9129 | TP53 | 7157 | Cell 145, 787-799, 2011 | |
| C14orf43 | 91748 | TP53 | 7157 | Cell 145, 787-799, 2011 | |
| DNTTIP1 | 116092 | TP53 | 7157 | Cell 145, 787-799, 2011 | |

**Supplementary Table 8**
