## supplementary table 9 for "A yeast BiFC-seq method for genome-wide interactome mapping"

| | No. of PPIs | Coexpression (SCC>0.5) | Ration of coexpression |
|---|---|---|---|
| negatome | 1152 | 131 | 0.113715 |
| P53 interactors | 96 | 20 | 0.208333 |
| Genome-wide interactors | 214 | 28 | 0.130841 |

**Supplementary Table 9**
